# Limb Specific Failure of Proliferation and Translation in the Mesenchyme Leads to Skeletal Defects in Diamond Blackfan Anemia

**DOI:** 10.1101/2022.01.14.476336

**Authors:** Jimmy Hom, Theodoros Karnavas, Emily Hartman, Julien Papoin, Yuefeng Tang, Brian M. Dulmovits, Mushran Khan, Hiren Patel, Jedediah Bondy, Morris Edelman, Renaud Touraine, Geneviève Chanoz-Poulard, Gregory Ottenberg, Robert Maynard, Douglas J. Adams, Raymond F. Robledo, Daniel A Grande, Philippe Marambaud, Betsy J Barnes, Sébastien Durand, Anupama Narla, Steven Ellis, Leonard I. Zon, Luanne L. Peters, Lydie Da Costa, Jeffrey M. Lipton, Cheryl L. Ackert-Bicknell, Lionel Blanc

## Abstract

Ribosomopathies are a class of disorders caused by defects in the structure or function of the ribosome and characterized by tissue-specific abnormalities. Diamond Blackfan anemia (DBA) arises from different mutations, predominantly in genes encoding ribosomal proteins (RPs). Apart from the anemia, skeletal defects are among the most common anomalies observed in patients with DBA, but they are virtually restricted to radial ray and other upper limb defects. What leads to these site-specific skeletal defects in DBA remains a mystery. Using a novel mouse model for RP haploinsufficiency, we observed specific, differential defects of the limbs. Using complementary *in vitro* and *in vivo* approaches, we demonstrate that reduced WNT signaling and subsequent increased β-catenin degradation in concert with increased expression of p53 contribute to mesenchymal lineage failure. We observed differential defects in the proliferation and differentiation of mesenchymal stem cells (MSCs) from the forelimb versus the hind limbs of the RP haploinsufficient mice that persisted after birth and were partially rescued by allelic reduction of *Trp53*. These defects are associated with a global decrease in protein translation in RP haploinsufficient MSCs, with the effect more pronounced in cells isolated from the forelimbs. Together these results demonstrate translational differences inherent to the MSC, explaining the site-specific skeletal defects observed in DBA.

## INTRODUCTION

Ribosomes are essential organelles involved in protein translation. They have long been thought to be so fundamental to cellular behavior that variations in the biosynthesis or function of these complex macromolecules were considered a biologic anathema. Thus, cellular activity was felt to be programmed at the DNA level and selected at the transcriptional level with the ribosome, an invariable machine, deciphering only the messages available to be translated. Recently, studies have highlighted the ribosome as a cell type-specific regulator of mRNA translation along with temporal regulation during development (Genuth & Barna, 2018). Thus, the complexity of the eukaryotic ribosome alone, consisting of four structural ribosomal RNAs and approximately 80 small and large subunit ribosomal proteins (RPs) offer an ample opportunity for modifications that permit both temporal and tissue specific regulation of translation (Gay et al., 2021; Shi & Barna, 2015). In particular, cell-specific ribosomes have been shown to be variable in their RP composition in different tissues and at different times in development (Shi & Barna, 2015).

Defects in RP expression or rRNA processing lead to a category of syndromes known as ribosomopathies (Narla & Ebert, 2010). Diamond Blackfan anemia (DBA) is one type of the known and characterized ribosomopathies that occur in humans (Da Costa et al., 2020). DBA is an inherited bone marrow failure syndrome, characterized by red cell aplasia, poor skeletal growth and congenital anomalies. Depending on the mutation, these can include craniofacial defects such as oral-facial clefts, vertebral fusions, scapular defects, radial ray anomalies and shortening of the long bones of the upper limbs. These patients also have a predisposition to cancer, notably osteogenic sarcoma (Vlachos et al., 2018). Despite these distinct defects, studies have mostly focused on the mechanisms leading to the red cell failure in these patients, and the mechanisms leading to skeletal defects and osteosarcoma in DBA are largely unknown. Most of the cases of DBA are due to a mutation in genes encoding a ribosomal protein involved in the ribosome assembly and the lack of which (Ulirsch et al., 2018) leads to the downstream cellular consequences of RP haploinsufficiency, including faulty translation and p53 activation (Da Costa et al., 2020).

Both the hematopoietic and mesenchymal lineages contribute to bone homeostasis (Calvi & Link, 2015; Yu & Scadden, 2016). While mesenchymal stem cells (MSCs) have the potential to differentiate into osteoblasts and chondrocytes, contributing to bone formation; hematopoietic stem cells (HSCs) can generate osteoclasts, essential for bone resorption. As mentioned above, patients with DBA have short stature. The presence of anemia together with treatments with corticosteroids leading to steroid induced bone loss and transfusion dependent iron overload may all contribute to poor linear growth (Sieff, 1993). For these reasons, the skeletal defects and the low bone mineral density (BMD) observed in DBA patients have not been well studied as a primary autonomous feature of the bone lineage but instead as consequences of anemia and standard treatments.

Recently, a study reported intra-uterine growth restriction in a stillborn fetus with a *de novo* mutation in *RPS19 (Da Costa et al., 2013)*. Here, we identify *RPS19* as a significant contributor of neonatal and adult skeletal development. By using complementary experimental systems including human induced Pluripotent Stem cells (iPSCs), mouse embryonic stem cell (mESCs) and a novel conditional mouse model for *Rps19* haploinsufficiency, we provide evidence that the targeted deletion of one allele of *Rps19* leads to disruptions in the mesenchymal lineage and that the bone defects seen in DBA are not solely secondary events caused by the standard of care treatments DBA patients. Specifically, reduced *Rps19* expression results in a limb-specific failure of MSC differentiation into the osteoblast lineage likely due to reduced global translation in these cells. Together, these results suggest a new pathogenic mechanism in bone development and maturation.

## RESULTS

### Skeletal Defects in a Patient with DBA

While numerous studies have reported skeletal defects in patients with a *RPL5* or *RPL11* mutation, these have not been as thoroughly investigated in patients with *RPS19* mutations. A recent case study highlighted growth defects in a stillborn male fetus delivered at ∼37 weeks of gestation with a *de novo* mutation in *RPS19* (Da Costa et al., 2013). As demonstrated by x-ray images, this fetus with DBA presented with shortened long bones of the upper limbs and absence of the deltoid tuberosity of the humerus (**Figure 1A**). To evaluate the cancellous bone architecture of this fetus, we used Masson’s trichrome and Safranin O staining on the rib bones and compared them to rib bones of a non-hematological control stillborn child at the same gestational age (**Figure 1B**). The cancellous bone or bony trabeculae appeared to show reduced connectivity of each trabecular element to adjacent trabeculae in the patient with DBA. We also noted a relatively thinner calcified matrix centrally within trabeculae and thicker, non-mineralized osteoid matrix on the surface of trabeculae, suggesting a defect in bone mineralization. These observations suggested involvement of the osteoblastic lineage in the skeletal defects of DBA.

**Figure 1.**
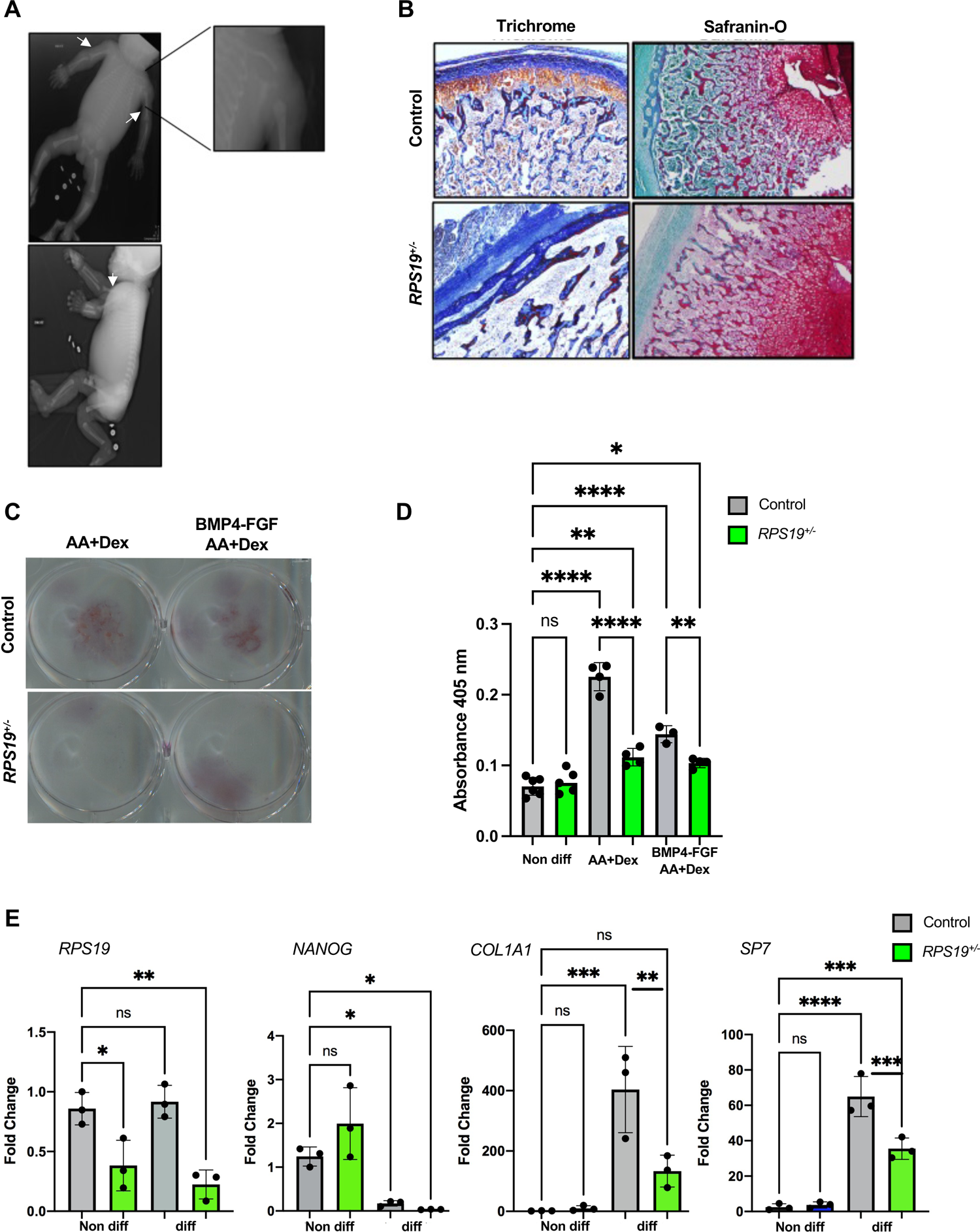
Clinical presentation of DBA caused by an Exon 2 mutation in *RPS19* and the phenotype of osteoblasts derived from osteoblast mediated by loss of one allele of *RPS19*. **A.** X-ray images of 37-week old male fetus heterozygous for a deletion in Exon 2 of *RPS19* that died *in utero.* White arrows denote short proximal upper extremities (inset image) with a posteroanterior view in the top image and a lateral view in the bottom image. **B.** Masson’s Trichrome and Safranin O staining of rib bones from the DBA patient and gestational age matched *RPS19*^+/+^ stillborn fetus. **C.** Whole six-well plate wells Alizarin Red staining of hiPS derived osteoblasts at day 30 of differentiation from *RPS19*^+/+^ and *RPS19*^+/-^ patients upon Ascorbic Acid+dexamethasone and FGF/Ascorbic Acid+dexamethasone induction. **D.** Quantification of the Alizarin Red staining of hiPS derived osteoblasts at day 30 of differentiation (n=3-6± SEM). **E.** Real time qPCR for mRNA levels of *RPS19*, the pluripotency marker *NANOG*, and two osteogenic markers (SP7 - Osterix and COL1A1 – Type I collagen) for undifferentiated *RPS19*^+/+^ and *RPS19*^+/-^ hiPSCs and respective cells differentiated to be osteoblasts. Gene expression was measured at day 30 of differentiation. Fold change relative to wt mESC and normalized for GAPDH. n=3, ± SEM.

### Mineralization by osteoblasts carrying mutations in *RPS19* derived from human iPSCs

Given the hypo-mineralization observed in the fetal bone and to acquire insights into the bone pathology of DBA, we differentiated iPSCs towards the osteoblast lineage (Doulatov et al., 2017). Specifically, these iPSCs were generated from fibroblasts from two individuals, both of whom were unrelated to the still born fetus. The patient with DBA carries a C280T mutation in the *RPS19* gene (*RPS19*^+/-^), and the second is a normal control with the native or wildtype allele for *RPS19* (Doulatov et al., 2017). We used two different approaches to differentiate these iPSCs, according to established protocols (Phillips et al., 2014). In both cases, we observed a decrease in mineralization 13 days after differentiation in the *RPS19*^+/-^ cells, although the difference was more pronounced when we differentiated cells in presence of ascorbic acid (AA) and dexamethasone (**Figure 1C&D**). The expression of RPS19 was reduced by half in the DBA patient-derived iPSCs (**Figure 1E**). Cells differentiated with a cocktail of dexamethasone and Fibroblast Growth Factor (FGF) had reduced expression of Osterix (*SP7*) and Type I Collagen (*COL1A1)*, which are markers of osteoblast commitment and maturation, respectively (**Figure 1E**). Together, these data suggest that mutations in *RPS19* can lead to skeletal defects in patients with DBA, independently of hematologic-related defects.

### *Rps19* Haploinsufficiency in Bone Progenitors Induces Defects in Osteoblastogenesis

To further confirm this finding in a different experimental cell model, we then used a mouse embryonic stem cell (mESC) model as, in our hands, this model has been highly reliable and reproducible to study ribosomal defects. Specifically, we previously reported that mESCs harboring a gene trap mutation in *Rps19* exhibited RP haploinsufficiency and polysome defects (Singh et al., 2014). Using a previously described method of *in vitro* differentiation (Kanke et al., 2014), we took advantage of this system to generate murine osteoblasts that were either wildtype or haploinsufficient for *Rps19* (*Rps19*^+/-^, **Figure 2A**). Accordingly, we observed a decrease of both *Rps19* transcript (**Figure 2B**) and RPS19 protein throughout differentiation (**Figure 2C&D**). By Day 12 of differentiation, we observed a robust mineralization in wildtype cells while in contrast, *Rps19*^+/-^ cells presented only sporadic nodule formation. At Day 23, we noticed a 50% decrease in alizarin red staining, consistent with a decreased mineralization (**Figure 2E-F**). At the molecular level, we assessed the expression of different markers of differentiation towards the mesoderm (**Figure 2G**) and osteoblastogenesis (**Figure 2H&I**) by qPCR. We noticed a mild but significant increase in *Nanog* mRNA at day 0 of culture and a slight reduction in *Brachyury* and *Mixl1* in the *Rps19*^+/-^ cells at day 8 and 5, respectively, therefore ruling out a major defect in the ability to engage into mesoderm differentiation (**Figure 2G**).

**Figure 2.**
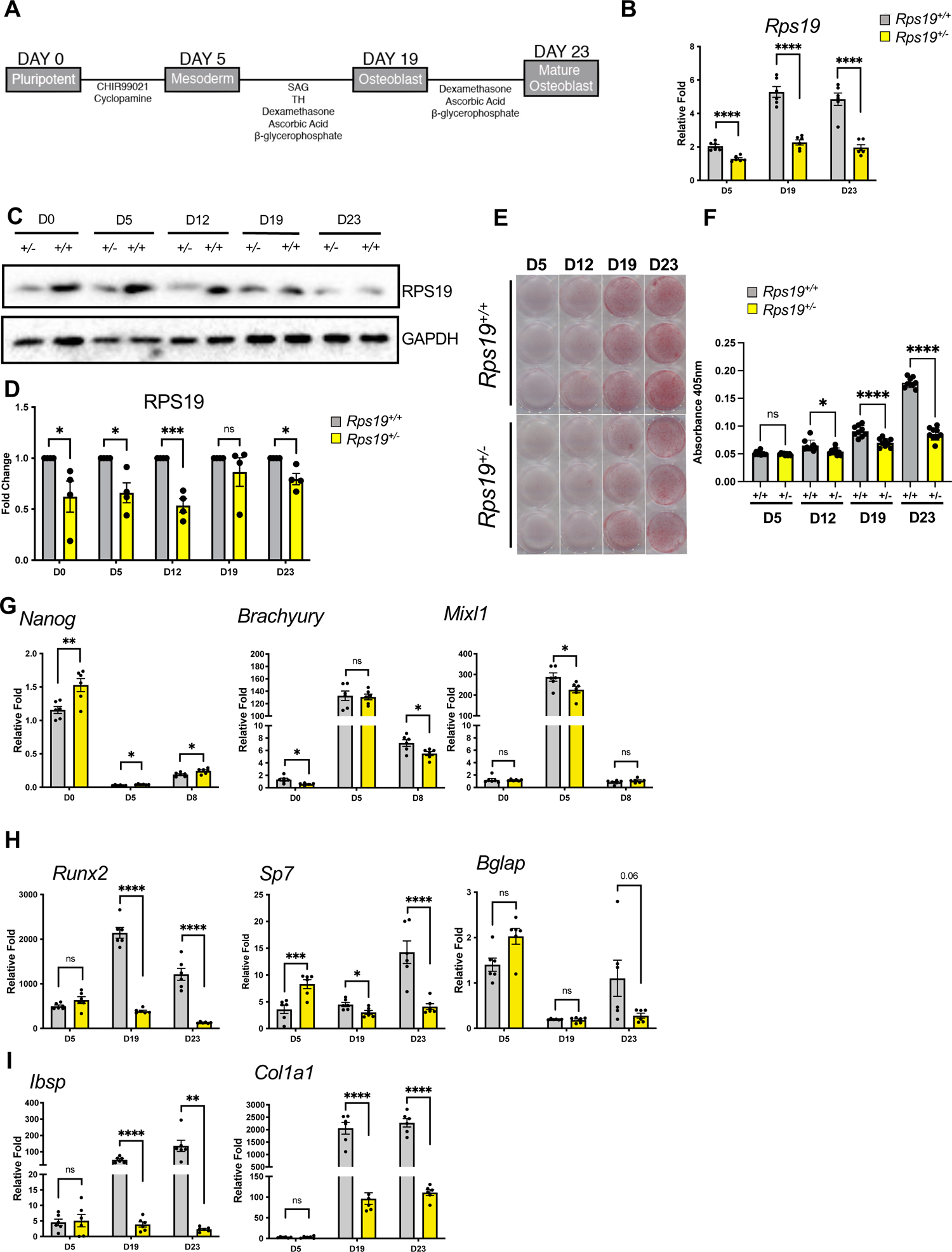
Loss of one allele of *Rps19* in mouse embryonic stem cells does not impact stemness but blunts the ability to differentiate these cells into osteoblasts. A. Experimental outline of stepwise *in vitro* osteogenic differentiation. B. Real time qPCR for *Rps19* mRNA levels at different time points of differentiation for *Rps19^+/+^* and *Rps19^+/-^* mESCs. Expression fold change relative to *Rps19^+/+^* mESC and normalized to *Gapdh*. n=6, ± SEM. C&D. Western blot and the quantification thereof from cell whole lysates for RPS19 content during mESC differentiation for *Rps19^+/+^* and *Rps19^+/-^* mESCs. Fold change relative to *Rps19^+/+^* mESC and normalized for GAPDH on a per time point basis. E. Alizarin S Red staining of whole 24-well plate wells during different four points of osteogenic differentiation of *Rps19^+/+^* and *Rps19^+/-^* mESC and the quantification of Alizarin S Red staining of the cells (n = 6-9 ± SEM). G-I. Real time qPCR for mRNA levels for genes denoting pluripotency (*Nanog*) and early mesodermal commitment (*Brachyury, Mixl1*) and markers of osteoblasts maturation *Rps19^+/+^* and *Rps19^+/-^* mESCs. n=6, ± SEM. *Runx2* and *Sp7* (Osterix) are key transcription factors for osteoblastogenesis, *Bglap* (Osteocalcin) is marker of mature osteoblasts and *Ibps* (Bone Sialoprotein) and *Col1a1* (Type I collagen) code for constituents of the extracellular matrix. Expression fold change relative to *Rps19^+/+^* mESC and normalized to *Gapdh*. n=6, ± SEM.

We then examined the expression of the two most critical osteoblast transcription factors, *Runx2* and *Sp7* (Osterix) across the 23-day culture period (**Figure 2H**). We noticed no difference in *Runx2* at D5, a time point reflecting the mesoderm stage of development, but curiously observed an increase in *Sp7* transcripts in the *Rps19*^+/-^ cells. However, at day 19 and 23, the expression of both transcription factors was substantially reduced in *Rps19*^+/-^ cells. Expression of *Bglap* (Osteocalcin), a marker of mature osteoblasts, was drastically reduced in the *Rps19*^+/-^ cells at day 23 (**Figure 2H**). We observed a similar pattern of reduced expression for two key bone extracellular matrix genes, *Ibsp* (Bone Sialoprotein) and *Col1a1* (Type I collagen, **Figure 2I**). Collectively, these data suggest that the loss of a single *Rps19* allele prevents bona fide osteoblast maturation by most likely affecting osteoblast formation and late differentiation steps rather than early engagement into mesoderm lineage differentiation. These results were consistent with observations made in human iPSCs from patients.

### Canonical WNT Signaling is Reduced in the *Rps19* Haploinsufficient mESC-derived Osteoblasts

To further explore the mechanism responsible for osteoblast maturation defects in *Rps19*^+/-^ cells, we examined the expression of the pro-osteoblastic genes by quantitative PCR arrays. mRNA levels were measured specifically at 23 days of culture to heighten the population of mature osteoblast-like cells. In addition to confirming the decrease in *Runx2, Col1a1* and *Col1a2 expression in Rps19*^+/-^ cells compared to *Rps19*^+/+^ cells, we also observed a decreased expression of *Itgav* and *Itga2*, which are implicated in osteoblast maturation (Olivares-Navarrete et al., 2011), as well as decreased expression of several members of the BMP/TGFB pathway (**Supplemental Fig 1**), further substantiating defects into osteoblast formation and maturation upon RPS19 depletion. Strikingly, the expression of several members of the BMP/TGFB pathway was reduced while we noticed an increased expression of *Tnfsf11* and *Itga*, which are both known to drive osteoblast-mediated induction of osteoclastogenesis (Ishii et al., 2009; Pederson et al., 2008).

Increased expression of P53 is a fundamental defect found in ribosomopathies, including DBA (Sieff et al., 2010), and is known to suppress WNT signaling (Kim et al., 2011). As both canonical and non-canonical WNT signaling pathways drive osteoblastogenesis and are associated with many skeletal dysplasia and developmental disorders (Wang et al., 2014), we examined the expression of a broad number of WNT signaling factors and targets using qPCR array. Interestingly, we observed a lower expression of the non-canonical pro-osteoblast *Wnt5A* gene, which is particularly interesting due to its association with Robinow syndrome (Person et al., 2010; Roifman et al., 2015) (**Figure 3A**). Indeed, Robinow syndrome patients exhibit craniofacial and limb length phenotypes, which, while not strictly identical to DBA, do display similar features to some extent. We also observed a decreased expression of the non-canonical WNT pathway Dvl3 gene (**Figure 3C**), the mutations of which cause an autosomal dominant Type III Robinow syndrome (White et al., 2016). Canonical WNT signaling is critical for many aspects for skeletal development and osteoblast function, including for osteoblast differentiation and bone matrix formation. Indeed, many loci identified by Genome Wide Association studies (GWAS) have linked canonical WNT signaling to low bone mineral density in humans ((Duan & Bonewald, 2016)). As WNT5A enhances canonical WNT signaling (**Figure 3B**) during osteoblast maturation (Okamoto et al., 2014) and has complex interactions with the canonical WNT receptor LRP6 during development (Gray et al., 2013), we then examined protein levels of this WNT receptor and the downstream mediator of canonical WNT signaling, β-catenin during osteoblast differentiation (**Figure 3C-D**). First, we observed reduced expression levels of both native and phosphorylated LRP6. Depending on its phosphorylation site, β-Catenin is either activated (phosphorylation on S552 and S675) or targeted for degradation through the ubiquitin/proteasome pathway (phosphorylation on S33/37/41, **Figure 3B**). We observed a decrease in S552/S675 phosphorylated β-Catenin from D12 to D23 of differentiation in *Rps19*^+/-^ cells while, in contrast, levels of S33/37/41 phosphorylated β-Catenin phosphorylation were strongly increased. Collectively, these results suggest that the activation of canonical WNT signaling is strongly diminished in *Rps19*^+/-^ cells (**Figure 3C&D**) and therefore provide strong evidence that loss of one allele of *Rps19* negatively impacts osteoblast maturation in the mesenchymal lineage. Curiously, this is associated with impaired canonical WNT signaling pathways, and given the known functions of WNTs in osteoblast maturation, these data implicate this pathways as being involved in this *Rps19*^+/-^ model of osteoblast maturation failure.

**Figure 3.**
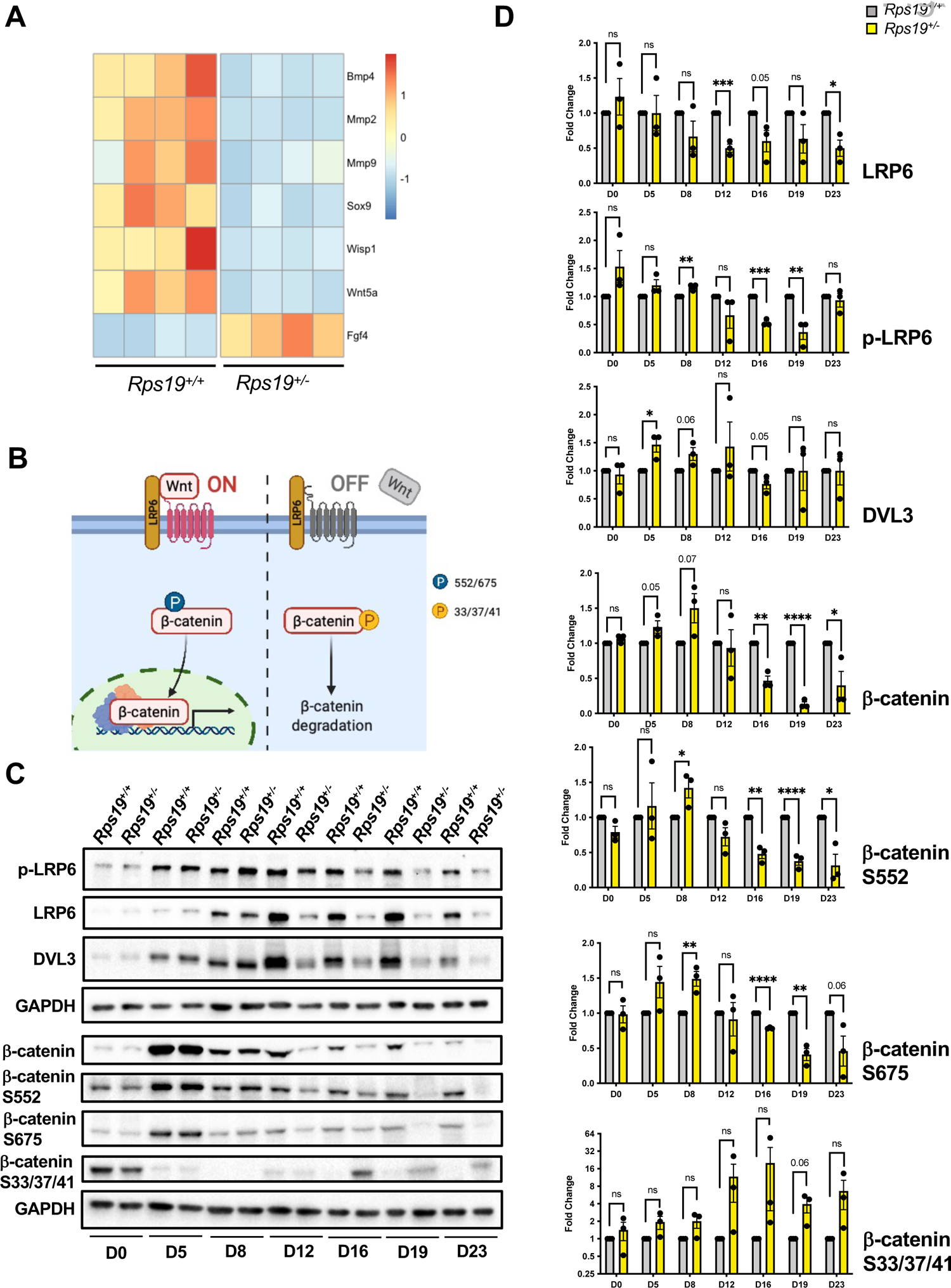
Characterization of canonical WNT signaling in osteoblasts derived from mESCs. **A.** Real Time PCR array quantification for select WNT pathway, WNT target genes associated with osteoblast function in *Rps19^+/+^* and *Rps19^+/-^* mESC derived osteoblasts on day 12 of differentiation (n=4, ± SEM). **B**. A cartoon representation of canonical WNT signaling and the phosphorylation mediated targeting of β-catenin for degradation (created with BioRender) **C.** Western blots from cell whole lysates for main components upstream of β-catenin in the canonical WNT signaling pathway (p-LRP6, LRP6, DVL3) and phosphorylated forms of β-catenin (S552, S675, S33/37/41) during mESC differentiation of *Rps19^+/+^* and *Rps19^+/-^* mESCs. **D**. Quantification of protein content for the same WNT pathway components at different time points of *Rps19^+/+^* and *Rps19^+/-^* mESC differentiation. Fold change relative to *Rps19^+/+^* mESC and normalized for GAPDH.

### Mice Lacking One Allele of *Rps19* in Mesenchymal Progenitor Cells Have Smaller Body Size, Limb Length Discrepancies and Cranial Defects at Birth

To dissect the role of *Rps19 in vivo*, we generated a conditional knockout allele for *Rps19* in which exon 2, containing the ATG translation start codon, was flanked by *loxP* sites (**Figure 4A**). Genomic PCR confirmed the presence of the *loxP* recombination sites (**Supplemental Figure 2**). Based on our data from mESCs (**Figures 2&3**), we hypothesized that *RPS19* reduction would impact skeleton development by affecting the early mesenchymal lineage. Therefore, we generated *Rps19^+/-^* mice where *Cre* expression is driven by the paired-related homeobox gene-1 (*Prrx1*) promoter beginning in the mesenchymal stem cell and continuing through the osteoblast precursor stage (Logan et al., 2002). The *Prrx1-Cre^+^, Rps19^+/lox^* animals were born with a normal Mendelian ratio (33 *Prrx1-Cre^+^, Rps19^+/lox^* live born of 64 total offspring) with a specific deletion of *Rps19* in both endochondral and intramembranous bone as well as in the femoral articular cartilage, as measured at both the genomic (**Figure 4B**) and at the protein level for the long bones (**Figure 4C&D**). In the femur, the expression levels of RPS19 were lowered by about 50% (**Figure 4D**) while levels in heart, liver and kidney remains identical to those of *Prrx1-Cre^+^; Rps19^+/+^* mice, as expected. *Prrx1-Cre^+^, Rps19^+/lox^* pups were of smaller size at birth (**Figure 4E**) and about 80% of the pups died by postnatal day 4 (P4), presumably due to their incapacity in feeding related to the limb defects, as it has been documented for other conditional models under a *Prrx1-cre* promoter (Lu et al., 2015). We did not observe *Prrx1-cre; Rps19^lox/lox^* animals, suggesting that *Rps19* is essential for the proper development of the mesenchymal lineage.

**Figure 4.**
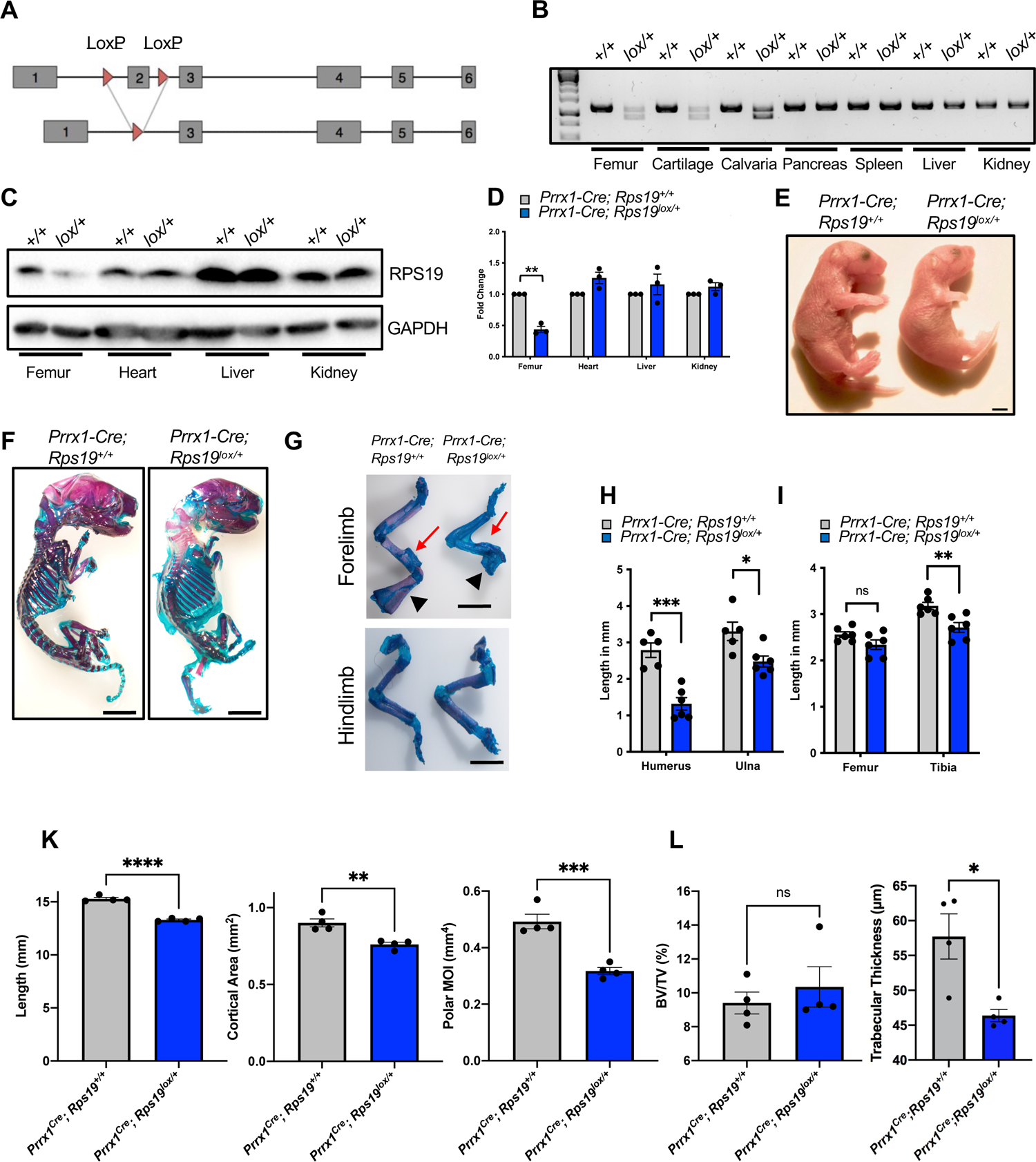
Mice heterozygous for and exon 2 deletion of *Rps19* in the mesenchymal lineage present with forelimb dysmorphology and growth plate defects. A. Genetic strategy for Cre-induced deletion of Exon 2 of *Rps19* under the promoter of *Prrx1*. B. Genomic PCR for specific recombination of *Rps19* allele in tissues of osteogenic lineage compared to other tissues in *Prrx1-Cre Rps19^+/+^* and *Prrx1-Cre Rps19 ^lox/+^* mice. C. Western blot for RPS19 from cell whole lysates of different tissues from neonatal *Prrx1-Cre Rps19^+/+^* and *Prrx1-Cre Rps19 ^lox/+^* mice. D. Quantification of RPS19 protein at different tissues from neonatal *Prrx1-Cre Rps19 ^lox/+^* mice. Fold change relative to *Prrx1-Cre Rps19^+/+^* mice and normalized for GAPDH. n=3, ± SEM. E. Whole body images of neonatal P0 *Prrx1-Cre Rps19^+/+^* and *Prrx1-Cre Rps19 ^lox/+^* mice. F. Whole mount Alizarin S Red and Alcian Blue staining of neonatal *Prrx1-Cre Rps19^+/+^* and *Prrx1-Cre Rps19 ^lox/+^* mice. Red staining denotes mineralized tissues and blue structures are cartilaginous. G. Alizarin S Red and Alcian Blue staining of forelimbs and hindlimbs of neonatal (P0) *Prrx1-Cre Rps19^+/+^* and *Prrx1-Cre Rps19 ^lox/+^* mice. The red arrow denotes the humoral tuberosity, and the black arrows point to the scapula. H-I: Quantification of length of forelimb bones (H) and hindlimb bones as noted (I) of neonatal (P0) *Prrx1-Cre Rps19^+/+^* and *Prrx1-Cre Rps19 ^lox/+^* mice. n=5, ± SEM. K-L. Quantitative measurements of femoral parameters derived by μCT imaging of 10-week-old male *Prrx1-Cre Rps19^+/+^* and *Prrx1-Cre Rps19 ^lox/+^*. (BV/TV=bone volume over total volume; MOI – moment of inertia) (n=4, ± SEM).

Alizarin red and Alcian blue staining of the whole skeleton and the forelimbs at P2 showed hypoplasia of the humerus and a striking agenesis of the scapula (**Figure 4F&G**). These defects were maintained 10-week old mice that survived (**Supplemental Figure 3A**). In addition, the deltoid tuberosity was missing in *Prrx1-Cre^+^, Rps19^+/lox^* (**Figure 4G**, red arrows). Unsurprisingly, this was associated with significant reduction in humoral and ulnar length (**Figure 4H**). At birth, the hindlimbs of *Prrx1-Cre^+^, Rps19^+/lox^* animals appeared less affected, but we did observe a slight but significant decrease in tibial length (**Figure 4H&I**). As the *Prrx1-Cre* is known to be active in selected craniofacial mesenchymal lineages, we also examined the calvaria in neonatal mice. Heterozygous mice presented with an increased opening of the anterior fontanelle due to an incomplete growth of the posterior frontal bone; but this defect resolved by day P21 (**Supplemental Figure 3B**). Consistent with patients with DBA harboring a mutation in *RPS19*, we did not observe any incidence of orofacial clefting. Together, these data suggest that the contribution of *Rps19* to skeletal development is site-specific with the most severe defects observed in the forelimbs.

### Loss of one allele of Rps19 impacted the cortical and trabecular bone of the femur

We then examined the impact of *Rps19* loss on the young adult cortical and trabecular bone compartments. While 10 week old mice are still rapidly growing, trabecular bone loss begins at or just before weaning (Glatt et al., 2007). We chose 10 weeks of age as the male mice are sexually mature, but not old enough to have experienced substantial maturity associated cancellous loss in the long bone. The femur was examined as this is the site typically examined in transgenic mouse models to look for cortical and trabecular bone defects (Rowe et al., 2018). The haploinsufficient femurs of 10-week old male mice were shorter than for the age matched wild type mice (**Figure 4K**). As expected, this resulted in a smaller cortical bone cross-sectional area at the mid-diaphysis and a highly significant reduction in polar 2nd moment of area (polar moment of inertia, POI). This latter measure suggests that the femurs of the haploinsufficient mice would be weaker in a twisting breaking configuration (**Figure 4K**). The ratio of the maximum diameter to the minimum diameter of the femur, a measure of roundness, was significantly smaller in the haploinsufficient femurs, likely contributing to the lower polar moment. As such, the bones of the haploinsufficient mice were not only anatomically smaller overall but also demonstrated a different overall morphology beyond the gross anatomical differences noted above (**Figure 4F&G**). While we saw no difference in the trabecular bone volume fraction (the ratio of trabecular bone volume to the total volume of the cancellous compartment), there was a significant difference in trabecular thickness (**Figure 4L**), which is reminiscent of the trabecular configuration we observed for the stillborn fetus. The *Prrx1*-cre is known not to be expressed in the axial skeleton (Durland et al., 2008) and therefore we could not determine if *Rps19* haploinsufficiency impacted the axial skeleton in our model.

### Dose-Dependent Activation of p53 Drives Skeletal Defects in *Rps19^+/-^* Mice

As mentioned above, p53 plays a key role in ribosomopathies. To determine if p53 was involved the defects observed in *Prrx1-cre; Rps19^lox/+^* animals, we used the mESC model of *Rps19* haploinsufficiency we generated above. While levels of *Trp53* mRNA remained unchanged during *in vitro* differentiation assays (**Figure 5A**), we observed an increase of activated (phosphorylated at S15) P53 levels beginning at D8 of culture and persisting into the mature osteoblast stage (**Figure 5B-D**), suggesting that p53-dependent apoptosis may play roles in *in vitro* osteoblast differentiation defects as well as *in vivo* skeletal defects observed in *Prrx1-cre; Rps19^+/lox^* mice. Therefore, we then hypothesized that reducing the levels of p53 could rescue the bone phenotype of the *Rps19^fl/+^* animals, as observed for the hematopoietic failure in other models of ribosomal haploinsufficiency. To test this hypothesis, we crossed the *Prrx1-Cre^+^; Rps19^fl/+^* mice with the *Trp53^fl/fl^* allele in order to generate mice lacking one or both copies of *Trp53* in the mesenchymal lineage. We observed a p53 dose-dependent rescue of with regards to whole body size at birth (**Figure 5E, Supplemental Figure 4**). This was also associated with *Trp53* dose-dependent resolution of the humoral and scapular morphology, and a complete correction of the cranial defects seen at birth (**Figure 5F**). These results are consistent with the overall observation that loss of *Rps19* impacts long bone morphology and development via the mesenchymal lineage and that p53 is involved in the DBA skeletal dysplasia, independently of its involvement in the hematopoietic lineage.

**Figure 5.**
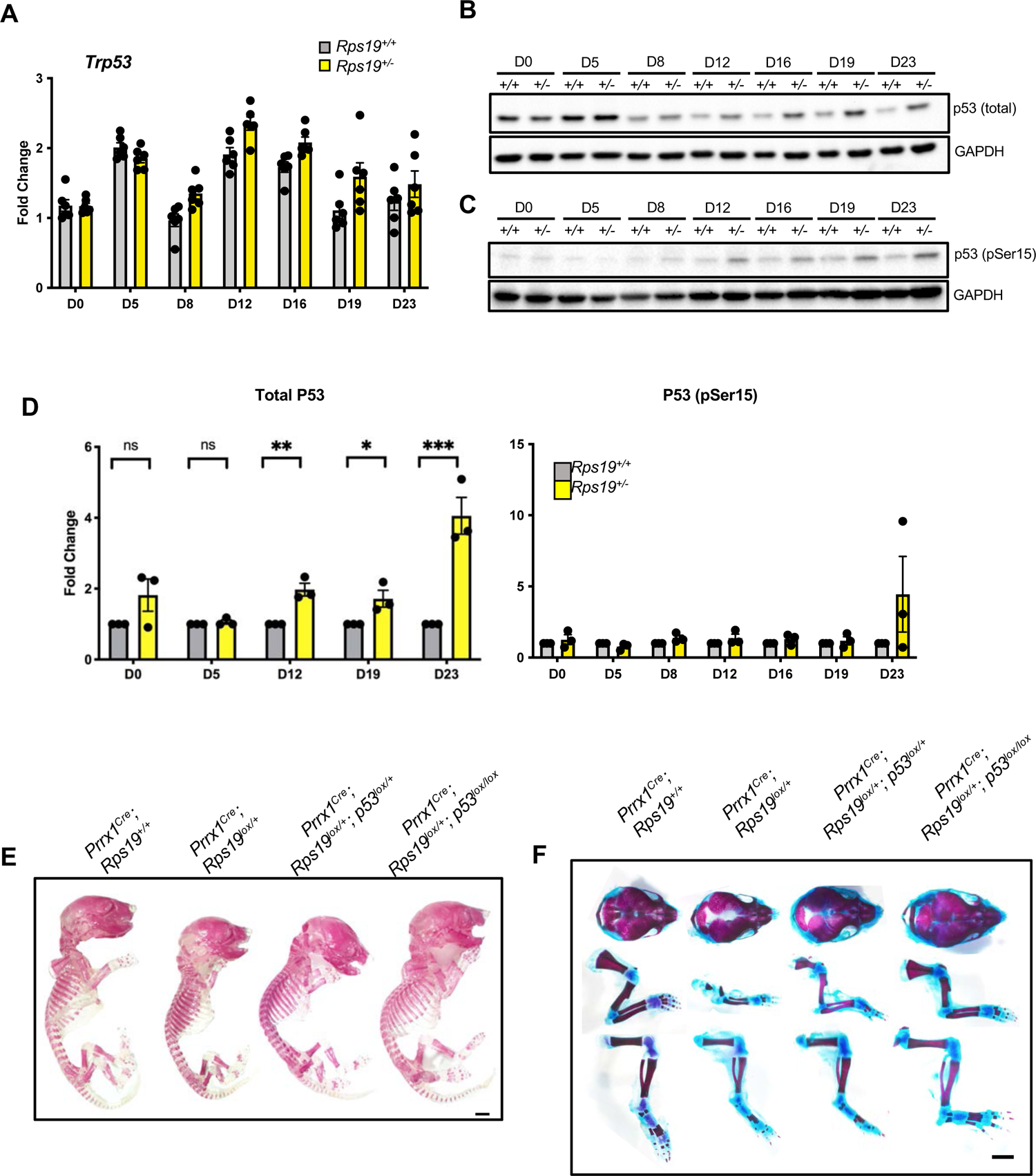
Allele dose dependent loss of *Trp53* rescues the skeletal phenotype seen with *Rps19* haploinsufficiency. **A.** Quantification of total P53 protein along osteogenic differentiation for *Rps19^+/+^* and *Rps19^+/-^* mESCs. Fold change relative to *Rps19^+/+^* mESC differentiated cells and normalized for GAPDH. n=3, ± SEM. **B-D.** Western blot and quantification for total TRP53 and pSer15 TRP53 during mESC differentiation. **E.** Alizarin S Red whole mount skeletal staining of neonatal (P0) *Prrx1-Cre Rps19^+/+^* and *Prrx1-Cre Rps19^lox/+^*, *Prrx1-Cre Rps19^lox/+^;Trp53^lox/+^*, *Prrx1-Cre Rps19 ^lox/+^;Trp53 ^lox/+^* mice. **E.** Alizarin S Red and Alcian Blue staining of calvaria, forelimbs and hindlimbs of neonatal (P0) *Prrx1-Cre Rps19^+/+^* and *Prrx1-Cre Rps19 ^lox/+^*, *Prrx1-Cre Rps19 ^lox/+^;Trp53 ^lox/+^*, *Prrx1-Cre Rps19 ^lox/+^;Trp53 ^lox/+^* mice.

### Forelimb-derived Mesenchymal Stem Cells from Rps19 Haploinsufficient Mice Fail to Differentiate due to defects in proliferation and translation

Our findings that the skeletal defects were more severe in the forelimbs than in the hindlimbs of *Rps19^lox/+^* animals suggested a unique developmental and or stem cell characteristic was evident in the forelimbs as this gene appears to be universally expressed in the MSC lineage. To address this question, we isolated Mesenchymal Stromal Cells (MSCs) from the forelimbs and hindlimbs of 3-4 week old mice irrespective of gender (**Figure 6** and **Supplemental Figure 5**) and differentiated them toward osteoblasts. We observed similar decreases in alkaline phosphatase activity in the *Rps19 ^lox/+^*-derived cultures, independent of the origin of the MSC (i.e. forelimbs or hindlimbs, **Figure 6A&B**). Strikingly, we noticed a dramatic and more pronounced reduction in mineralization in *Rps19^lox/+^*cultures derived from the forelimbs when compared to the hindlimbs as measured by alizarin red staining (**Figure 6C&D**). These results suggest that MSC derived from the humeri are more affected by *Rps19* haploinsufficiency and those isolated from the femur.

**Figure 6.**
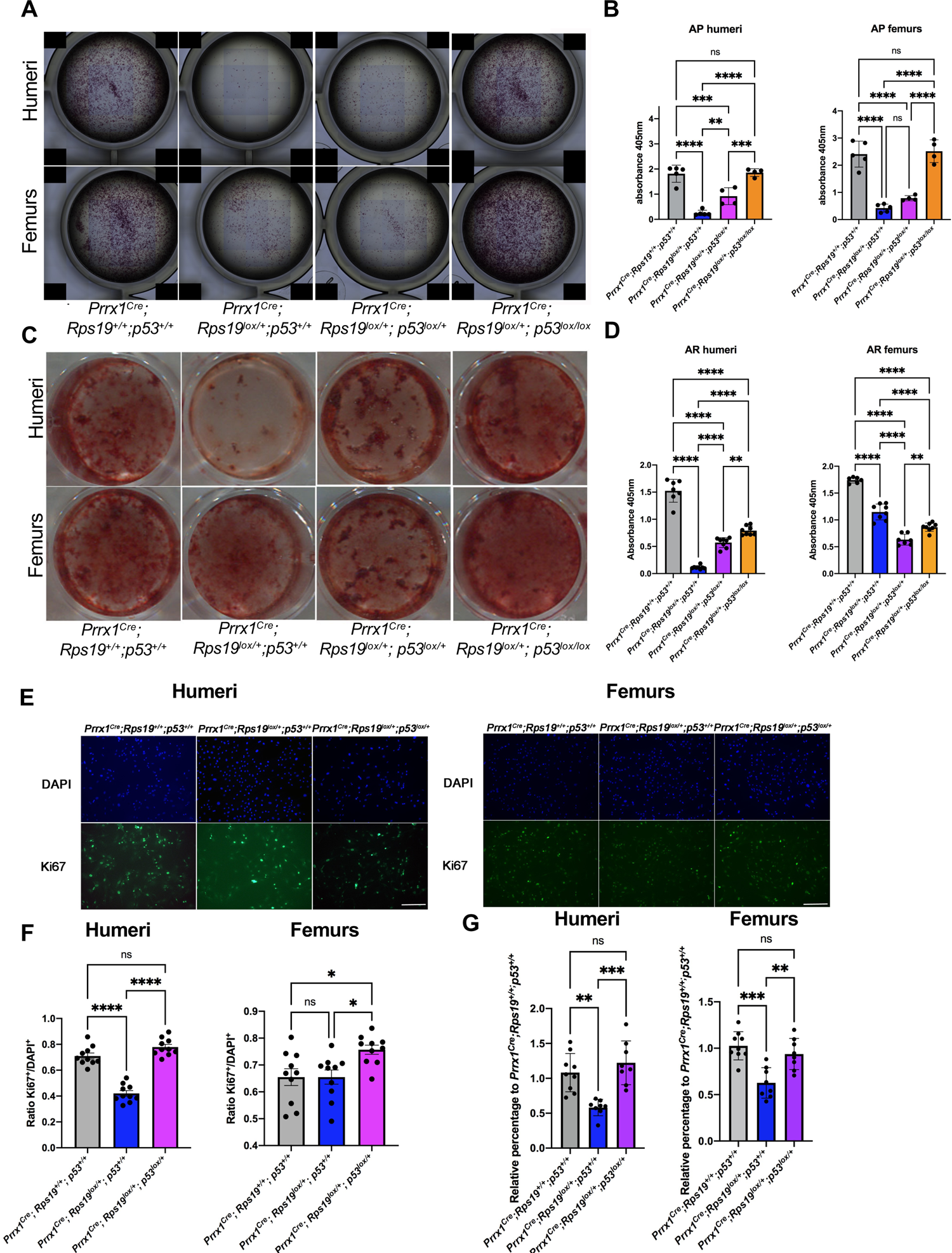
Characterization of the osteoblastic potential of mesenchymal stem cells (MSCs) isolated from the humeri and femurs of 3 weeks old *Rps19^+/-^* mice. **A.** Alkaline phosphatase staining of whole 24-well plates wells of MSC at passage 2. **B.** Quantification of alkaline phosphatase staining of whole 24-well plates wells of MSCs P2. (n=5, ± SEM). The data presented is sex aggregated data. Sex segregated data is presented in Supplementary Figure 5. **C.** Alizarin S Red staining of whole 24-well plates wells of MSCs derived osteoblasts at day 14 of differentiation from humeri and femurs of *Prrx1-Cre Rps19^+/+^*, *Prrx1-Cre Rps19^lox^*^/+^, *Prrx1-Cre Rps19^lox^*^/+^;*Trp53^lox^*^/+^, *Prrx1-Cre Rps19^lox^*^/+^;*Trp53^lox^*^/+^ mice. **D.** Quantification of Alizarin Red S staining of whole 24-well plates wells of MSCs derived osteoblasts at day 14 of differentiation. (n=7, ± SEM). The data presented is sex aggregated data. Sex segregated data is presented in Supplementary Figure 5. **E.** Staining for the proliferation marker ki67 in MSCs from the humeri (left) and the femur (right) at passage 2 (scale bar=200μm). **F.** Quantification of ki67 staining for MSC the humeri and femurs. (n=10, ± SEM). **G**. Flow cytometry and quantification for OPP incorporation the humeral and femoral derived MSCs of *Prrx1-Cre*; *Rps19*^+/+^, *Prrx1-Cre; Rps19*^lox/+^, *Prrx1-Cre*; *Rps19*^lox/+^*p53l^ox^*^/+^ 3-week old mice. (n=10, ± SEM). The data presented is sex aggregated data. Sex segregated data is presented in Supplementary Figure 5.

To get further insights into the mechanism responsible for the specific decrease in humoral-derived MSCs differentiation, we first evaluated the proliferative capacity of the isolated MSCs. Using anti-Ki67 immunofluorescence microscopy, we observed a specific decrease in proliferation of MSCs isolated from the humeri, while proliferation was unaffected in the femoral-derived MSCs (**Figure 6E&F**). Importantly, we observed a dose-dependent rescue of the phenotype in MSC derived from *Rps19^lox/+^* with p53 deleted (**Figure 6A-F**). Together, our results suggest that the more pronounced defect exhibited by the humeri may be due, at least in part, to the specific failure of proliferation of the humoral-derived MSCs.

A fundamental part of bone formation is the apposition of the proteaceous extracellular matrix component of bone. Therefore, we evaluated protein translation efficiency of the MSCs isolated from *Rps19^lox/+^* animals. To do so, we incubated MSCs with O-propargyl-Puromycin (OPP) and then measured the fluorescence intensity by flow cytometry. We observed a global decrease in protein synthesis in the *Rps19^lox/+^* total cell population independent of the bone origin (**Figure 6G**). However, we observed a specificity in translation efficiency depending on the source of the MSC; the humoral-derived MSCs being more susceptible to *Rps19* haploinsufficiency in terms of decrease in translation (**Figure 6G, left panel**). Interestingly, the defect in translation was rescued by the removal of p53, as very recently documented in an *Rps6* mutant mouse (Tiu et al., 2021), however this work was only conducted using cells from a single bone type. Altogether, these results suggest that the more severe defects observed in the humeri compared to the femur from *Rps19* haploinsufficient animals may be due to specific decreased proliferation and protein synthesis in humoral-derived MSCs.

## DISCUSSION

In this study, we established that *Rps19* is required for normal skeletal patterning and growth. Using multiple different models in which *Rps19* levels were decreased by ∼ 50% in cells recapitulating the mesenchymal lineage, we directly associate defects in a ribosomal protein with a failure of normal bone formation and with improper limb growth and skeletal formation. The models we have used are complementary and specifically allow for comprehensive characterization of the mesenchymal failure due to *Rps19* haploinsufficiency. We specifically show that loss of *Rps19* impacts osteoblast maturation, resulting in reduced collective osteoblast function due to defective WNT signaling. Using our conditional *Prrx1-Cre,Rps19^fl/+^* model, we established that *Rps19* is essential for both forelimb and hindlimb formation and growth, albeit to a different degree. This is associated with a bone specific (humeri vs femur) reduction in marrow derived mesenchymal stem cells (MSCs) and reduced protein translation by these cells. Further, we show that loss of p53 protein can rescue the long bone phenotype, including the MSC number, but can only partially correct for the protein translation defect.

Numerous studies have focused on the erythroid defects associated with DBA (Da Costa et al., 2020). Due to the obvious obstacles inherent in studying skeletal development and growth in patients much less effort has been undertaken to characterize DBA-associated congenital anomalies and skeletal defects. With regards to skeletal defects, this study is the first to recapitulate in the mouse the phenotype observed in patients and to demonstrate that the origin of the skeletal defect is a function of how mesenchymal cells function at different anatomic sites.

The stillborn fetus with a nonsense mutation in exon 2 of *RPS19* presenting with short upper limb extremities suggests that the role of *RPS19* in limb development is conserved during mammalian evolution. This study also demonstrates a pathogenic role for *RPS19* in determining the bone phenotype outside of the hematopoietic lineage, eliminating a singular role for the myeloid derived osteoclast. The skeletal system is essential for blood formation and niches within the bone marrow microenvironment that regulate hematopoietic stem cell fate specification (Teti & Teitelbaum, 2019). However, the *Prrx-Cre Rps19^fl/+^* mice do not present with anemia (**Supplemental Figure 6**), precluding an extrinsic role of the mesenchyme to the anemia observed in DBA and vice a versa.

We considered multiple possible scenarios that could explain the skeletal phenotypes we see in our mouse model and in DBA caused by *RPS19* mutations in general, recognizing the distinction between skeletal development and mature bone growth. As the mesenchyme matures, there is an increased demand for protein translation during development and more specifically by the chondrocyte and then osteoblast. In the first scenario we considered, with reduced RPS19, and the presumed reduction in ribosome function, the MSCs simply cannot keep up with needed protein synthesis, leading to skeletal defects and reduced bone. This scenario is supported by the reduced protein synthesis we see in the skeletally derived MSCs. Further, a major transcription factor involved in osteoblastogenesis is RUNX2, which was decreased in both the mESC and iPSC-derived osteoblasts. Loss of *Runx2* globally results in failure to form osteoblasts (Ducy et al., 1997). These same cells have decreased transcript levels of the gene *Sp7* (Osterix), compared to wildtype cells. This reduction in *Sp7* is not unexpected given the significant reduction in *Runx2* (Nakashima et al., 2002). Through a direct binding interaction, p53 prevents Osterix from binding to its Sp1/GC-rich sites and from interacting with the pro osteoblastogenesis transcription factor *Dlx5*, which in turn leads to the decreased production of the bone matrix constituents COL1A1 and Bone Sialoprotein (IBSP) (Artigas et al., 2017). This, in addition to the decreased translation, likely contribute to the reduced osteoblast mediated matrix mineralization we observed. However, this scenario cannot explain why the phenotype of DBA is so specific with regards to which anatomic features and physiological systems are impacted as there is a ubiquitous need by all MSC cells to translate mRNA into proteins.

In a second scenario, we conjectured that the ribosome concentration model (Danilova & Gazda, 2015; Gabut et al., 2020; Mills & Green, 2017) could explain some of the skeletal defects observed in DBA. Indeed, studies have demonstrated that in erythroid precursors, decreased translation of GATA-1 led to the anemia and can be rescued by overexpressing GATA-1 (Ludwig et al., 2014). In osteoblast progenitors and precursors, the same mechanism could exist, albeit targeting a different transcription factor. This would affect the MSC pool distribution leading to a selective feed forward effect through development. This latter scenario could explain why defects in DBA impact only certain tissues in discrete ways during development and is supported by the observation by other groups that there is more than one population of mesenchyme and more than one type of terminal MSC such as the osteoblast (Ambrosi et al., 2021). Further, anatomic dependent effects on different bone populations and the subsequent skeletal phenotype are common in global knockout transgenic mice (Rowe et al., 2018). This would explain our observation of the striking difference in the number MSCs derived from the humeri compared to the femur.

The phenotype seen in DBA is also a function of which signaling pathways are impacted by the loss of RPS19 and when they are affected. The lack of impact of *Rps19* reduction on the mesodermal transition from pluripotency in mES yet a profound impact on subsequent pre-osteoblast differentiation suggests that pathways driving MSC commitment are mechanistically the drivers of the skeletal dysplasia of DBA. There are numerous pathways involved in mesenchymal commitment and maturation, including the Bone Morphogenic Protein (BMP), Hox, Hedgehog, Fibroblast Growth Factor (FGF) and Sox pathways (Mortus et al., 2014). Mutations in many of these pathways impact limb development. Critically, the skeletal dysplasia found during limb development in our model was rescued by reduction of p53. It has been long appreciated that in ribosomopathies that p53 was elevated in part due to nucleolar stress due to defective protein translational machinery (Fumagalli & Thomas, 2011). Countless studies have demonstrated the critical roles for both canonical and non-canonical WNT signaling in skeletal morphogenesis, chondrogenesis, osteoblast maturation and general bone physiology (Duan & Bonewald, 2016). However, more recently it has been shown that loss of β-catenin, the downstream effector of canonical WNT signaling in MSCs, drives p53 expression and that reduction of p53 in the absence of β-catenin lifts the p53-mediated blockade on osteoblast maturation (Zhou et al., 2021). While the interactions between p53 and the WNT signaling pathway depend on the cellular context, studies have reported that activation of p53 leads to the down regulation of β-catenin (Kim et al., 2011). Together this suggests a complex loop between Trp53 and β-catenin. In our mES model, we see a reduction of the active form of β-catenin, an increase in the form of β-catenin targeted for degradation and coincident increase in p53. It cannot be determined from our data if the rise in p53 protein comes before the loss of β-catenin or if a loss of β-catenin contributes to the rise in p53, but our data does ultimately suggest that the intersection between p53 and WNT signaling plays a role in the skeletal dysplasia in DBA.

Finally, we observed a differential limb phenotype in our mouse model with forelimb displaying more severe developmental abnormalities than hindlimb. A possible explanation could be provided by the timing in the onset of *Prrx1* promoter activation, as *Prrx1* expression begins a day earlier in the forelimb at E9.5 (Logan et al., 2002). However, patients with DBA also have more severe defects in the radial ray and upper extremities although these patients have germline autosomal mutations impacting all cells starting from conception. This would suggest that the phenotype of our mouse is not a function of the timing of *Rps19* excision by the *cre* driver we chose, but rather due to a differential impact of a loss of *Rps19* in the mesenchyme. In conclusion, our study offers a novel explanation for the skeletal dysplasia phenotype associated with DBA.

## Material and Methods

### Human Subjects

All human studies were conducted in accordance with the world medical association declaration of Helsinki. The fetal tissues were collected with informed consent of the parents and under the supervision of the Institutional Review Board from the Hospital of St. Etienne, France. The fibroblasts used to make the iPSCs, which have been previously described (Doulatov et al., 2017), were obtained after informed consent of the Institutional Review Board of Boston Children’s Hospital, Boston, MA, USA.

### Transgenic Mice

All animal procedures were performed according to protocols reviewed and approved by The Feinstein Institutes for Medical Research Institutional Animal Care and Use Committee (IACUC). Where possible, data from public repositories was used to reduce the number of animals used in this study. The forms of euthanasia that were used were done so in accordance with the guidance and recommendations provided by the Panel on Euthanasia of the American Veterinary Medical Association.

*Prrx1*-*Cre* (JAX #005584) and *Trp53*^fl/fl^ (JAX #008462), were purchased from the Jackson Laboratory. A systemic loss of the *Rps19* gene in mice is not viable (Matsson et al., 2004). Therefore, we generated a floxed allele that would allow for tissue specific deletion of Exon 2, a critical exon in the mouse gene. Specifically, mice carry the *Rps19^fl^* allele were via CRISPR/cas9 endonuclease mediated genome editing of the *Rps19* locus using sgRNA sequences (GGGTGGACTGGCGACGAGCA, GGACTGGCGACGAGCAGGGT, TCTTTTCTGAATTGGGCCTA, AAAGGAAGCATGGTCACCGT). To ensure a consistent genetic background between the floxed allele and all other needed alleles for this study, mice carrying the *Rps19^fl^* allele were backcrossed at least 10 times with inbred a C57BL/6J mice. To generate mice lacking one allele of *Rsp19* in the mesenchymal lineage, male mice carrying *Prrx1*-*Cre* were bred to *Rps19^fl/+^* mice. This unidirectional cross was used to avoid the known potential for germline allele excision when *Prrx1*-*Cre* is passed from the dam (Logan et al., 2002). Mice carrying the *Trp53^fl^* allele were first crossed with mice carrying the *Rps19^fl^* allele and consequently *Rps19^fl/fl^ Trp53^fl/+^* and, *Rps19^fl/fl^ Trp53^fl/fl^*, *Rps19^fl/+^ Trp53^fl/fl^* female mice were crossed with *Prrx1 – Cre or Prrx1 –Cre Trp53^fl/+^* male mice. For all experiments, at least three replicates per genotype were analyzed. Mice were genotyped by PCR and the sequences for the genotyping primers to detect loxP sequences and *Cre*-mediated recombination are listed in **Supplemental Table** 1. The ages and genders of the mice used are listed for each experiment.

### Cell culture

#### Human induced pluripotent stem (hiPS) cell culture maintenance and differentiation

Wildtype (Wt) iPSCs cell lines were used along with hiPS cell lines from DBA patients carrying a nonsense mutation C280T as described in (Doulatov et al., 2017). Both wild type and mutant hiPS cell lines came from human skin fibroblasts reprogrammed with episomal method as previously described (Doulatov et al., 2017). iPS cells were cultured in tissue culture 10-cm dishes, prep-coated with hESCs qualified Matrigel (Fisher Scientific, 08774552) for 1hr at 37°C. The cells were cultured in mTESR1 medium (StemCell Technologies, 85857), medium was replenished daily and were passaged every 5-6 days with Dispase I (StemCell Technologies, 07923) in small cell clumps. iPSCs were frozen in 40% mESC qualified FBS (GIBCO, 16141061), 50% mTESR1 medium and 10% DMSO.

For iPSCs osteogenic differentiation, cells were plated in 6-well tissue culture plates that were pre–coated with Matrigel (Corning, 354234) one day before day 0 of differentiation. For osteogenic differentiation of iPSCs, we used the protocol previously described (Phillips et al., 2014) with the following modifications: cells were cultured in basal medium containing Knockout DMEM (GIBCO, 10829018), 10% FBS (Cytiva, SH30071.03), 1% penicillin-streyptomycin (Thermo Scientific, 15140122), 0.1 mM non-essential amino acids (Thermo Fisher Scientific, 11140050), 2 mM glutamax (Thermo Scientific, 11140050), either with the addition just prior to use of 0.01μM dexamethasone (Sigma-Aldrich, D4902), and 1μM ascorbic acid (AA) phosphate (Sigma-Aldrich, A92902) for 30 days or with the addition prior to use of 6ng/ml bFGF (R&D, 3718-FB-01M) for first 7 days (days 0-7), followed by addition of 0.01μM dexamethasone and 1μM ascorbic acid phosphate for last 23 days (days 8-30). Medium was replenished every 3 days.

#### Mouse Embryonic Stem Cell (mESC) culture maintenance and differentiation

The *Rps19* mutant mESC line YHC074 (Mutant Mouse Regional Resource Center, RRID:MMRRC_012362-UCD) was used along with parental wild type mouse ESC line E14Tg2a.4 at passages P5-8. mESCs were cultured in feeder free conditions with 5% CO2 at 37°C on 0.1% gelatin (StemCell Technologies, 7903) coated plates in 2i medium supplemented with 2% mES certified FBS (Gibco 16141061), 1000U/mL leukemia inhibitory factor (ESGRO LIF; Millipore, ESG1107), 1uM PD0325901 (Cayman, 13034), and 3μM CHIR99021 (Cayman, 13122) as previously described (Kanke et al., 2014). 2i basal medium consisted of 1:1 DMEM high glucose/F12 (GIBCO, 11330-057) and neurobasal medium (Gibco, A3582901), 1% penicillin-streyptomycin (Thermo Scientific, 15140122), 0.05% BSA fraction V (Sigma, A9576), 0.1 mM non-essential amino acids (Thermo Fisher Scientific, 11140050), 2 mM glutamax (Thermo Scientific, 11140050), N2 supplement (Thermo Scientific, #15140122), B27 supplement (Thermo), 0.055μM β-mercaptoethanol (Sigma-Alrich, 21985023). LIF, PD0325901, and CHIR99021 were supplemented to the media prior to use. mESCs were split using diluted 1:3 Accutase/PBS mixture and plated at 20,000 cells/cm2 for passaging. Cells were passaged every 2 days and medium was replenished daily. mESCs were frozen using 90% mES-FBS and 10% DMSO.

For mESC osteogenic differentiation, 24-well tissue culture plates were precoated using Matrigel (Corning, 354234) for 1hr at 37 °C and 15,000 cells/cm2 were plated 24 hours before day 0 of differentiation in complete maintenance medium. Osteoblast differentiation were performed for 23 days in 2i basal medium without LIF, serum and PD0325901 as previously described (Kanke et al., 2014) with the following modifications: Mesodermal induction was performed for days 0-5 using 15μM CHIR99021 and 5μM cyclopamine (Cayman, 11321). Pre-osteoblast differentiation was then induced for days 6-19 in 1μM SAG (Cayman, 11914) and 1μM helioxanthin analogue TH (Tokyo Chemical Industry, M3085) with the ostegenic inductive factors 0.1μM dexamethasone (Sigma-Aldrich, D4902), 50μg/mL ascorbic acid phosphate (Sigma-Aldrich, A92902), and 2mM β-glycerophosphate (Sigma-Aldrich, G9422). Osteoblast maturation was performed for an additional 4 days without SAG and TH and included dexamethasone, ascorbic acid phosphate, and β-glycerophosphate. PBS wash was performed on day 4 and cell media was changed daily.

#### Mesenchymal stem cell (MSC) isolation, cell culture and differentiation

Mesenchymal stem cells were isolated from limbs of 3-4 weeks old mice of both sexes, according to (Zhu et al., 2010). Bone chips were plated in 12-well and 6-well plates in MEM-A medium (GIBCO, 12561-056) supplemented with 10% FBS (Cytiva, SH30071.03), 1% penicillin-streyptomycin (Thermo Scientific, 15140122) and cultured for 5-7 days at P0 before been passaged with Tryspin - EDTA 0.05% (Thermo Scientific, 25300054). Cells were passaged every 4-5 days and until p8. Medium was replenished every 2 days. For differentiation of mesenchymal stem cells towards osteogenic, lineage cells of P2-5 were differentiated according to published instructions and protocols (Zhu et al., 2010). Cells were plated at a cell density of 5×10^3^ cells/cm^2^ in 24-well plates with MEM-A (GIBCO, 12561-056) supplemented with 10% (vol/vol) FBS (Cytiva, SH30071.03), 1% penicillin-streyptomycin (Thermo Scientific, 15140122), 10^-7^ M dexamethasone, 10mM β-glycerol phosphate and 10μM ascorbate – 2 – phosphate in a total volume of 500μl/well. Cells cultured in MEM-A with 10%FBS were used as control. Medium was changed every other day and cultures were maintained until day 21.

### RNA isolation, RT-qPCR and Gene Expression qPCR array

Total RNA from all types of cultured osteoblasts described herein was collected using RLT Qiagen RNAeasy mini/micro kit cell lysis buffer and further purified using RNAeasy mini (Qiagen, 74134) or micro (Qiagen, 74034) kit. To generate the cDNA, 1μg of RNA was reverse-transcribed using SuperScript VILO Master Mix (Invitrogen, 11766050) with ezDNase according to manufacturer’s instructions. Generated cDNA was diluted two to fivefold and qRT-PCR was performed using 2X Power SYBR Green PCR Master Mix (Applied Biosystems, 100029283) on a QuantStudio5 384-well PCR system. Biological replicates were run in technical triplicates. The housekeeping gene *Gapdh* was used as an internal control for normalization of expression values. Primer sequences are provided in **Supplemental Table 2**. Standard PCR cycling of 95 °C for 15 sec and 60 °C for 30 sec were used for 40 cycles, preceded by a 95 °C denaturation for 1min and followed by a Met-Curve between 60-95 °C. For qPCR array analyses, 1μg of RNA was reverse transcribed using RT2 First strand kit (Qiagen, 330404) and cDNA was used on the mouse osteogenesis (Qiagen, PAMM 026ZR) and WNT signaling targets PCR array (Qiagen, PAMM 243ZE-4) according to manufacturer’s instructions. Results were analyzed with the RT2 Profiler PCR Array Data Analysis.

### Western blot Analysis

Cultured osteoblasts of the types described above were lysed in 1X RIPA buffer (Thermo Scientific, 89900) supplemented with protease inhibitor cocktail (Bimake, B14002), phosphatase inhibitor cocktail III (Research Products International, P52104-1), vortexed at maximum speed and centrifuged at 4°C at 16,000 g for 10 min. Supernatant was diluted 1:1 with 2X Laemmli sample buffer (BioRad, 1610737) under reducing conditions and boiled for 5 minutes. Samples were separated via sodium dodecyl sulfate polyacrylamide gel electrophoresis (SDS-PAGE) for 1.5 hour at 100V and transferred to nitrocellulose membranes for 1 hr at 90V. Membranes were blocked with 5% (wt/vol) milk in 1x Tris-Buffered Saline and 0.1% (vol/vol) Tween20 unless specified otherwise by antibody manufacturer. Primary antibodies were incubated overnight at 4°C. List of primary antibodies used is provided in **Supplemental Table 3**. Membranes were washed and incubated with horseradish peroxidase (HRP)-conjugated secondary antibodies for 1 hour at RT. Bands were detected using enhanced chemiluminescence (ECL) (Thermo Scientific, PI32106). Quantifications were obtained using ImageJ software (NIH).

### Alizarin Red staining and quantification

Cultured and differentiated osteoblasts were fixed in 4% PFA/PBS for 15 minutes at RT, washed with 3x PBS at RT and stained with alizarin red S (Sigma Aldrich, A5533) 1% in ddH2O, pH 4.3 at 30 minutes at room temperature on a rocking platform. Whole stained wells were visualized for higher magnification images with EVOS M5000 Imaging System and scanned using an EPSON scanner. For quantitative assessment of alizarin red staining, fixed wells were washed with PBS 3x and completely dried before an equivalent volume to the culture media volume of 5% perchloric acid was added in all wells and placed on a rocking platform for 5 minutes at RT. Supernatants were collected and were quantified using Synergy Neo2 (Biotek) spectrophotometer at 405 nm using Alizarin Red standards of 0.0625, 0.125, 0.5, 1, 2 and 4mM Alizarin Red S solution in 5% perchloric acid.

### Alkaline phosphatase (AP) staining and quantification

Cultured osteoblasts were fixed with 4% PFA/PBS for 5 minutes at room temperature and stained with EMD Millipore staining kit (#SCR004) before imaged as whole wells using EVOS M5000 Imaging System. For quantification of AP activity, the Fisher Scientific (AS, #72146) kit was used according to manufacturer’s instructions.

### Ki67 Immunofluorescence

MSCs at P2 and calvaria osteoblasts at P0 were cultured on gelatin coated coverslips and were fixed with 4% PFA/PBS for 15 minutes at 4 °C and blocked for 1hr at room temperature with blocking buffer (0.1M PBS, 0.15% Glycine, 0.1% TritonX, 2mg/ml BSA). Wells were incubated with primary antibody against ki67 (Abcam, Ab16667) over night at 4°C at dilution of 1:250. Cells were washed 3x with PBST (0.1M PBS, 0.1% TritonX) at room temperature and then incubated for 1 hour at room temperature with a secondary Alexa Fluor goat anti - rabbit antibody at a concentration of 1:2000. Cells were washed again 3x with PBST at room temperature and stained with DAPI mounting medium (Sigma F6057). Coverslips with stained cells were visualized with EVOS M5000 Imaging System.

### μCT analysis

Femurs were dissected of soft tissues and fixed in 70% ethanol at 4°C from 10-week old male mice. Cortical morphometry of the femoral mid-diaphysis, and trabecular morphometry within the metaphyseal region of distal femurs, were quantified using X-ray micro-computed tomography (μCT50, Scanco Medical AG, Bassersdorf, Switzerland). Femurs were imaged in 70% ETOH at 70 kVp (200 μA), employing 2000 cone beam projections per revolution at an integration time of 500 msec. Three-dimensional images were reconstructed at ∼10 μm resolution using standard convolution back projection algorithms with Shepp and Logan filtering and rendered at a discrete voxel density of 1,000,000 voxels/mm^3^ (isometric 10 μm voxels). Bone mineral was calibrated to a discrete-step hydroxyapatite phantom and segmented from marrow and soft tissue in conjunction with a constrained Gaussian filter to reduce noise, applying mineral density thresholds of 700 and 500 mg HA/cm^3^ for cortical and trabecular bone, respectively. Anatomical boundaries of the diaphyseal cortex and the internal ROI within the trabecular compartment were identified using automated routines. Cortical morphometry was averaged within a mid-diaphyseal span of ∼0.7 mm (5% of femur length) to obtain measures of total area (Tt.Ar), marrow area (Ma.Ar), 2^nd^ moment of area (I), which is often referred to as Polar Moment of Inertia and the minimum and maximum dimensions of the mid diaphysis. Cortical morphometry was averaged within a mid-diaphyseal span of ∼0.7 mm (5% of femur length) to obtain measures of total area (Tt.Ar), cortical area (Ct.Ar), marrow area (Ma.Ar), polar Moment of Inertia (MOI), and the minimum and maximum dimensions of the mid diaphysis. Volumetric regions for trabecular bone analysis were selected within the endosteal borders to include the secondary spongiosa of femoral metaphyses located ∼0.7 mm (∼7% of length) from the growth plate and extending ∼1 mm proximally, scaling for bone length. Trabecular morphometric parameters were measured without imposing a presumed structural model to obtain direct measures of trabecular volume fraction (BV/TV), thickness (Tb.Th) and number (Tb.N, not shown).

### Whole Mount Skeletal Staining

Whole skeletal preparations for cartilaginous and mineralized skeletal elements performed as described (Rigueur & Lyons, 2014). Skeletal structures of newborn mice (P0) or 3-week old mice (P21) were dissected to remove the skin and fat and stained with Alizarin Red (Sigma-Aldrich A5533) for visualization of the mineralized elements and Alcian Blue (Sigma-Aldrich, B8438) to see the cartilaginous structures.

### Bone Immunofluoresence Microscopy and Histologic Staining

Femurs and humeri bones were dissected and fixed in 4% paraformaldehyde (PFA) at 4°C overnight. After washing in water, samples were then decalcified for 10 days using 14% (w/v) EDTA (pH 7.4), embedded in paraffin, and sectioned onto slides at 5μm. Hematoxylin and Eosin (H&E), Masson’s Trichrome,) staining were performed according to manufacturer’s instructions. The sections were then counterstained with 0.1% Fast Green. Bright field imaging was performed on an EVOS M5000 Imaging System. Data are representative of at least three independent experiments.

### Measurement of Global protein synthesis

For measuring of global protein synthesis with incorporation of O-Propargyl-Puromycin (OPP), mesenchymal stem cells (MSCs) at passage 1-2 isolated from 3-week old mice were labeled with 10μM OPP (MedChem Express, HY-15680) in MEM-A for 1hr. Subsequently the cells were washed with 1x PBS and trypsinised. Cells were washed with 3ml ice-cold PBS 0.1% BSA and proceeded with flow protocol as described in respective section.

### Flow Cytometry for protein synthesis measurement

Mesenchymal Stem Cells (MSCs) at passage 1-2 isolated from 3 weeks mice were trypsinised, counted and resuspended at a concentration of 10^7^/ml in ice – cold PBS 0.1% BSA. Cells were incubated with a list of antibodies (**Supplementary Table 4**) for 30’ on ice protected from light. Following antibodies incubation, cells were washed 1x with 3ml ice – cold PBS 0.1% BSA and then fixed with 4% PFA in PBS for 15’ at room temperature protected from light. Cells were washed with 3ml ice - cold PBS 0.1% BSA and resuspended in 100μl ClickIT kit (Thermo Fisher Scientific C10269) permeabilization buffer and incubated fro 15’ at room temperature before adding 400μl of the Plus reaction buffer of ClickIT kit, containing Alexa Fluor 555 Picolyl Azide dye (Thermo Fisher C10642), according to manufacturer’s instructions. Mix was incubated for 30’ at room temperature protected from light. Following the incubation, cells were washed 1x with 3ml permeabilization buffer, centrifuged and resuspended in 200μl of PBS 0.1% BSA and stained for 7-AAD (BD –Pharmingen 559925) for 10’ at RT protected from light before being analyzed in BD LSRFortessa Cell Analyzer. For fluorophore compensation OneComp eBeads from Invitrogen (01-1111-42) were used by mixing one drop of beads with 1μl of each antibody used in our panel while having only beads as control. The mix was incubated for 10’ at RT protected from light. 1ml of PBS 0.1% BSA and the mix was centrifuged and resuspended in 300μl PBS 0.1% BSA before analysis. For flow data analysis, Flowjo 10.5.3 software was utilized.

## Statistics

Statistical analysis was performed using Prism 8 (GraphPad Software). Unless otherwise specified, unpaired, two-tailed Student’s t test (single test) or one-way ANOVA (multiple comparisons) with a Tukey’s Post Hoc test were used to evaluate statistical significance (defined as *P<0.05*). All results in bar graphs are mean value plus SEM.

### Rigor and Reproducibility

Each experiment was conducted at least three independent times. All of the data are reported individually in each figure. Biological replicates represent individual animals while technical replicates are defined as repeated measurement on the same sample. No outliers were identified, no data were excluded.

## Acknowledgments

This work was supported by a World Allied St. Baldrick’s Foundation scholarship (to L.B.), a grant from the DBA foundation and DBA Canada (to L.B.), the National Institutes of Health 13U01HL134812 (to L.I.Z., J.M.L and L.B.) and P30CA034196 (to The Jackson Laboratory).

## Authorship

J.H., T.K., L.L.P., L.D.C., J.M.L., C.A-B. and L.B. designed the research; J.H., T.K., E.H., J.P., Y.T., B.M.D., M.K., H.P., J.B., G.O., R.M., D.J.A., R.F.R. performed the experiments; M.E., R.T. and G.C-P. contributed to the analyses of the skeletal defects in the still born patient; J.H., T.K., E.H., J.P., Y.T., B.M.D., M.K., H.P., J.B., G.O., R.M., D.J.A., R.F.R., D.A.G., P.M., B.J.B., S.D., A.N., S.E., L.I.Z., L.L.P., L.D.C., J.M.L., C.A-B. and L.B. analyzed and interpreted the data; J.H., C.A-B. and L.B. cowrote the manuscript; T.K., P.M., B.J.B., S.D., A.N., S.E., L.I.Z., L.L.P., D.J.A., and L.D.C. edited the manuscript; and all authors read and commented on the final manuscript.

## Supplemental Figure Legends

**Supplemental Figure 1:**
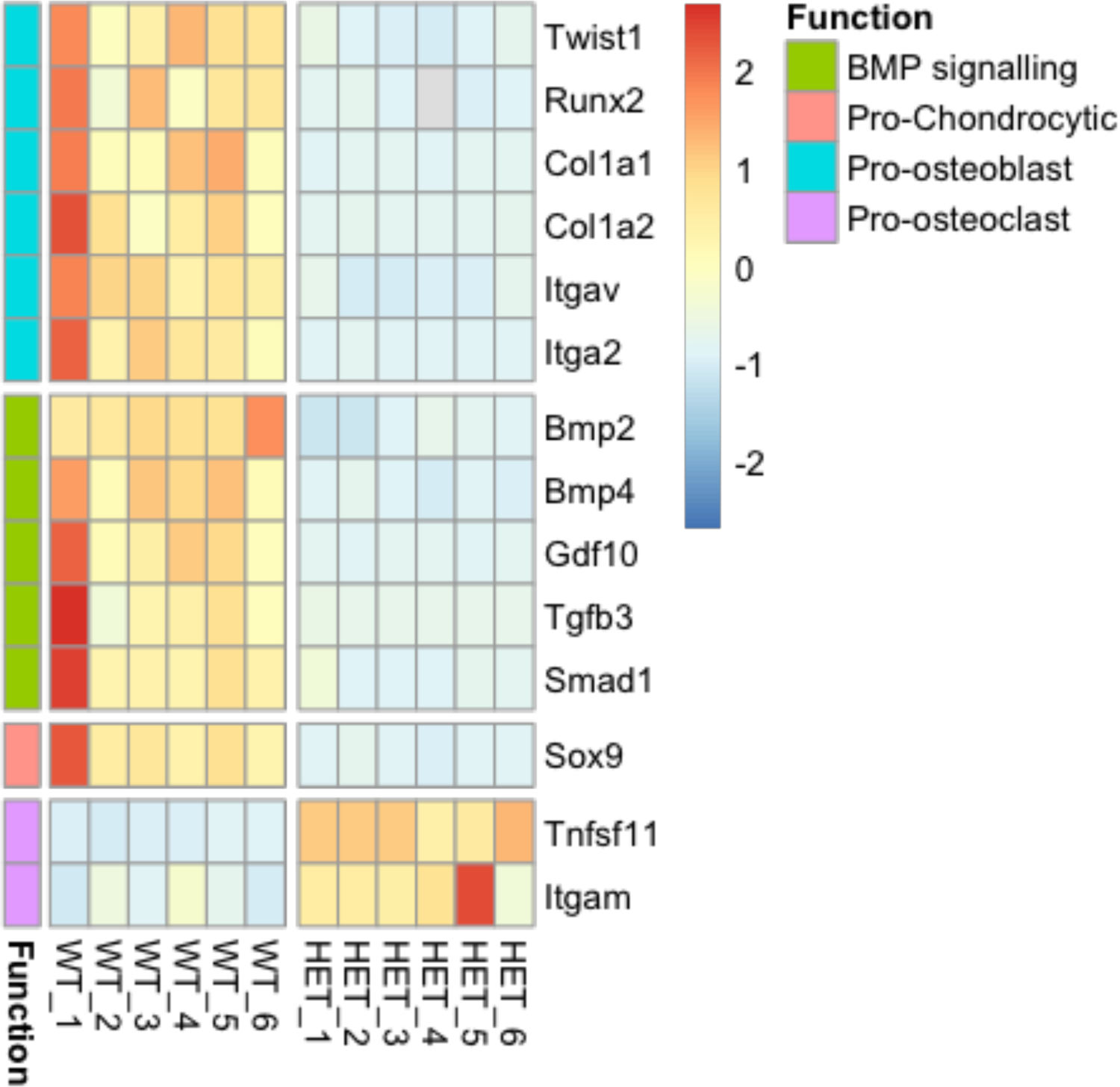
BMP signaling and osteo/chondro gene expression profile of *Rps19^+/-^* mESCs. Real Time quantitative PCR array for BMP signaling, pro – chondrocytic, pro – osteoblast and pro – osteoclast genes in *Rps19^+/+^* and *Rps19^+/-^* mEScs at day 23 of osteogenic differentiation. n=6

**Supplemental Figure 2:**
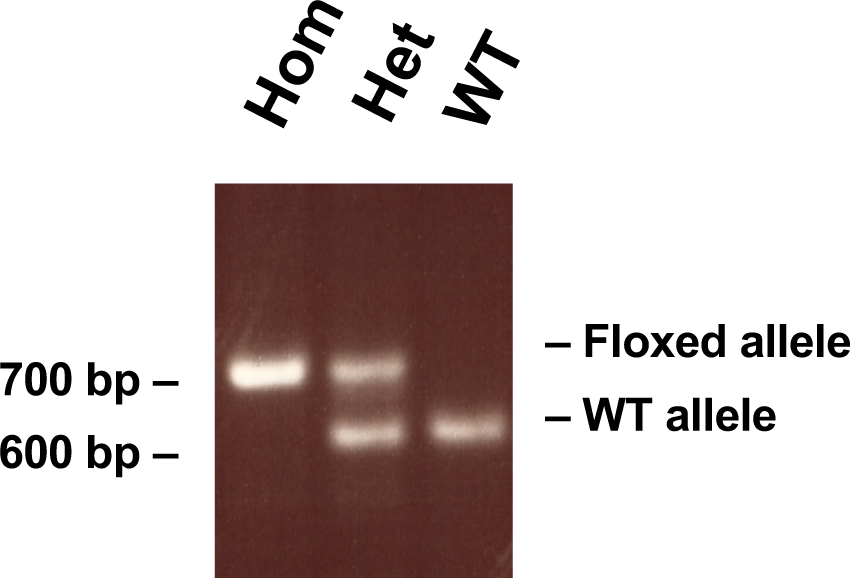
*Rps19^lox/+^* genotyping result. PCR reaction products for *Rps19^+/+^, Rps19^lox/+^* and *Rps19^lox/lox^* mice in 1.2% agarose gel. Product sizes: wt allele: 670bp, floxed allele: 760 bp.

**Supplemental Figure 3.**
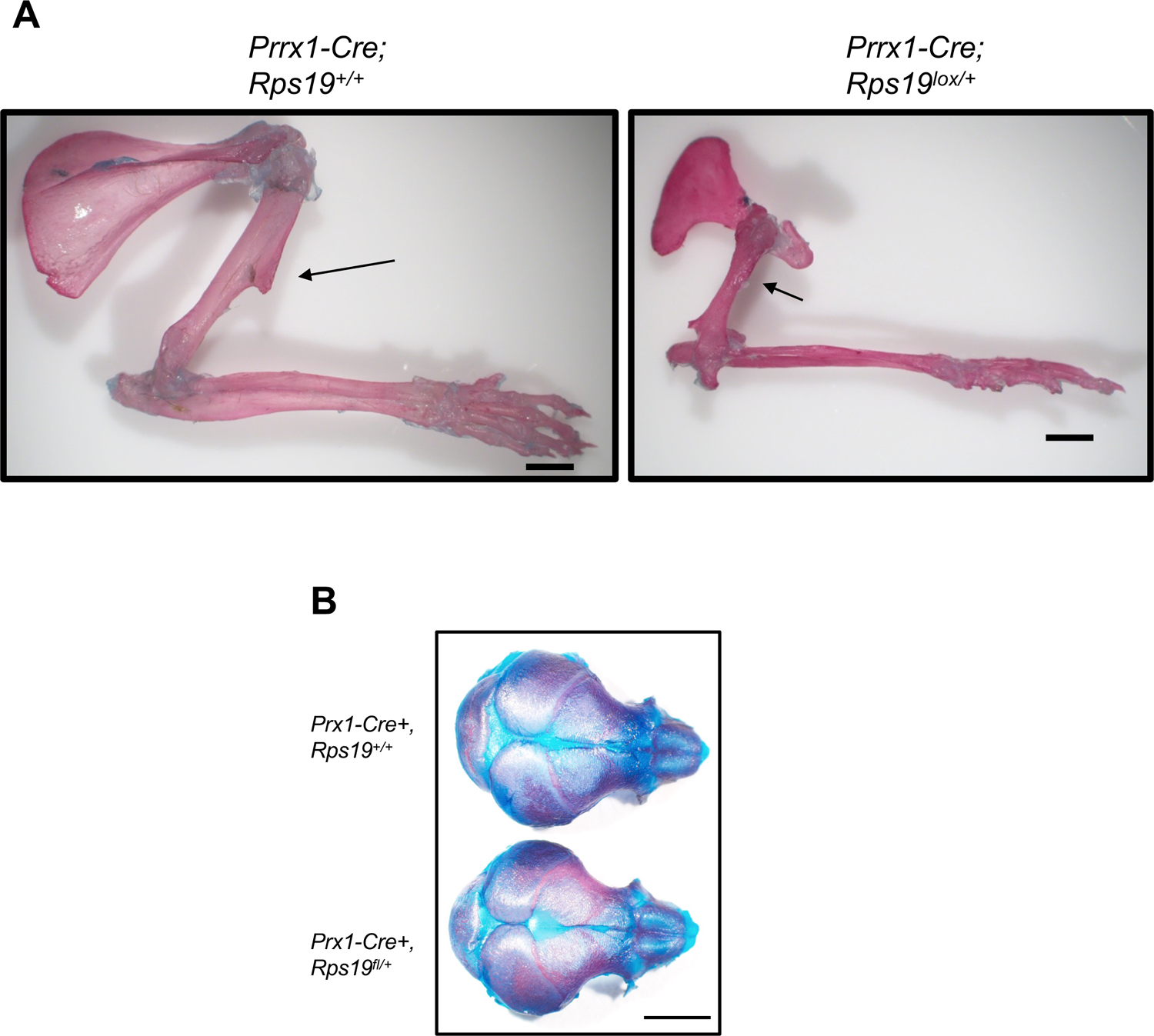
Skeletal defects of *Prrx1 – Cre; Rps19^lox/+^* mice. (A) Alizarin S red and alcian blue staining of forelimbs from 10 - week old male *Prrx1 – Cre; Rps19^+/+^* and *Prrx1 – Cre; Rps19^lox/+^* mice. Black arrows denote the deltoid tuberosity of the humerus. (B) Alizarin S red and alcian blue staining of P0 calvaria from *Prrx1 – Cre; Rps19^+/+^* and *Prrx1 – Cre; Rps19^lox/+^* mice. (Scale bar:2mm).

**Supplemental Figure 4.**
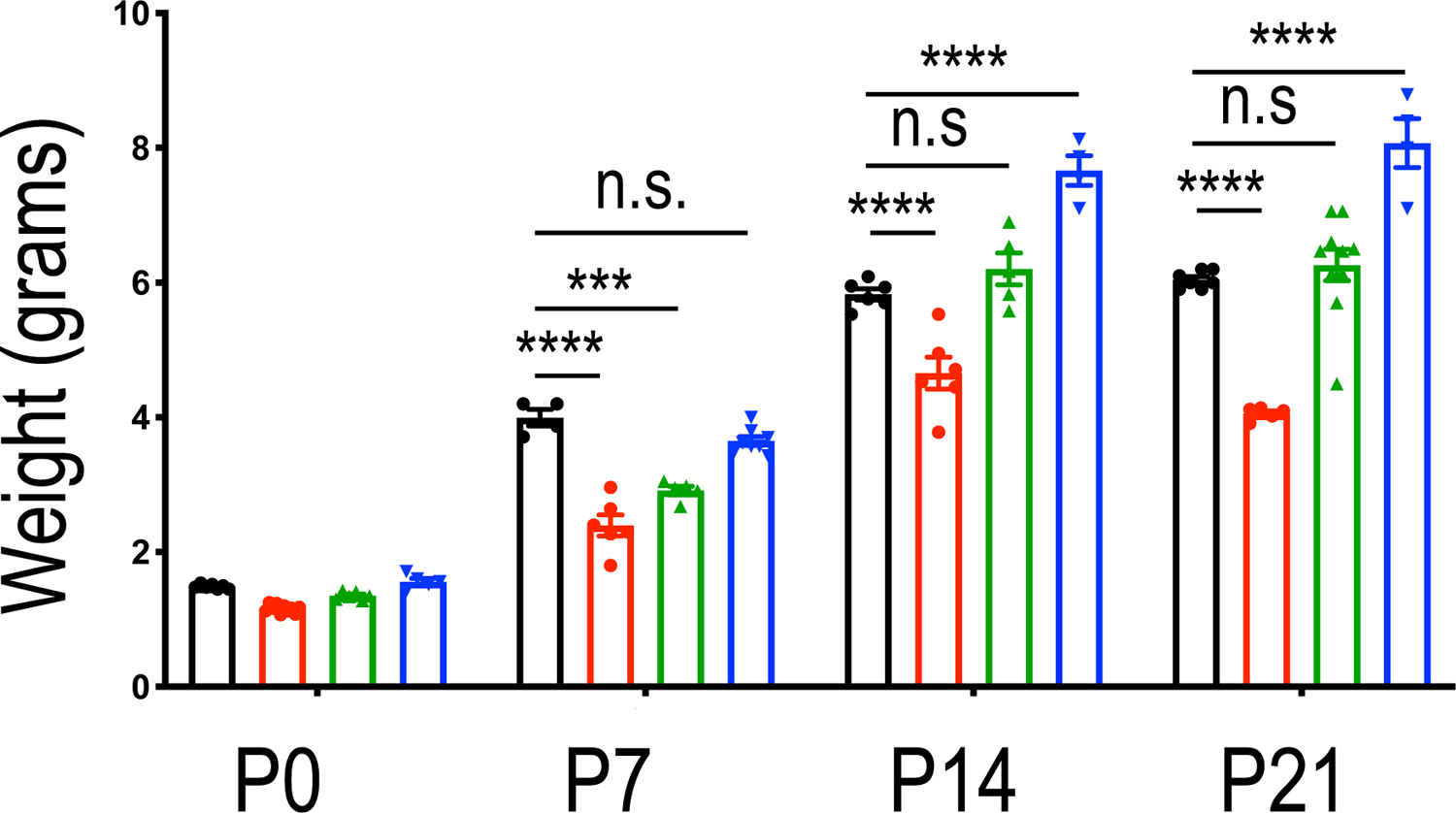
Weight phenotype of Prrx1-Cre; Rps19^lox/+^ mice. Weight measurements in grams of Prrx1-Cre; Rps19^+/+^ (black), Prrx1-Cre; Rps19^lox/+^ (red), Prrx1-Cre; Rps19^lox/+^p53^lox/+^ (green) and Prrx1-Cre; Rps19^lox/+^p53^lox/lox^ (blue) of P0, P7, P14 and P21.

**Supplemental Figure 5.**
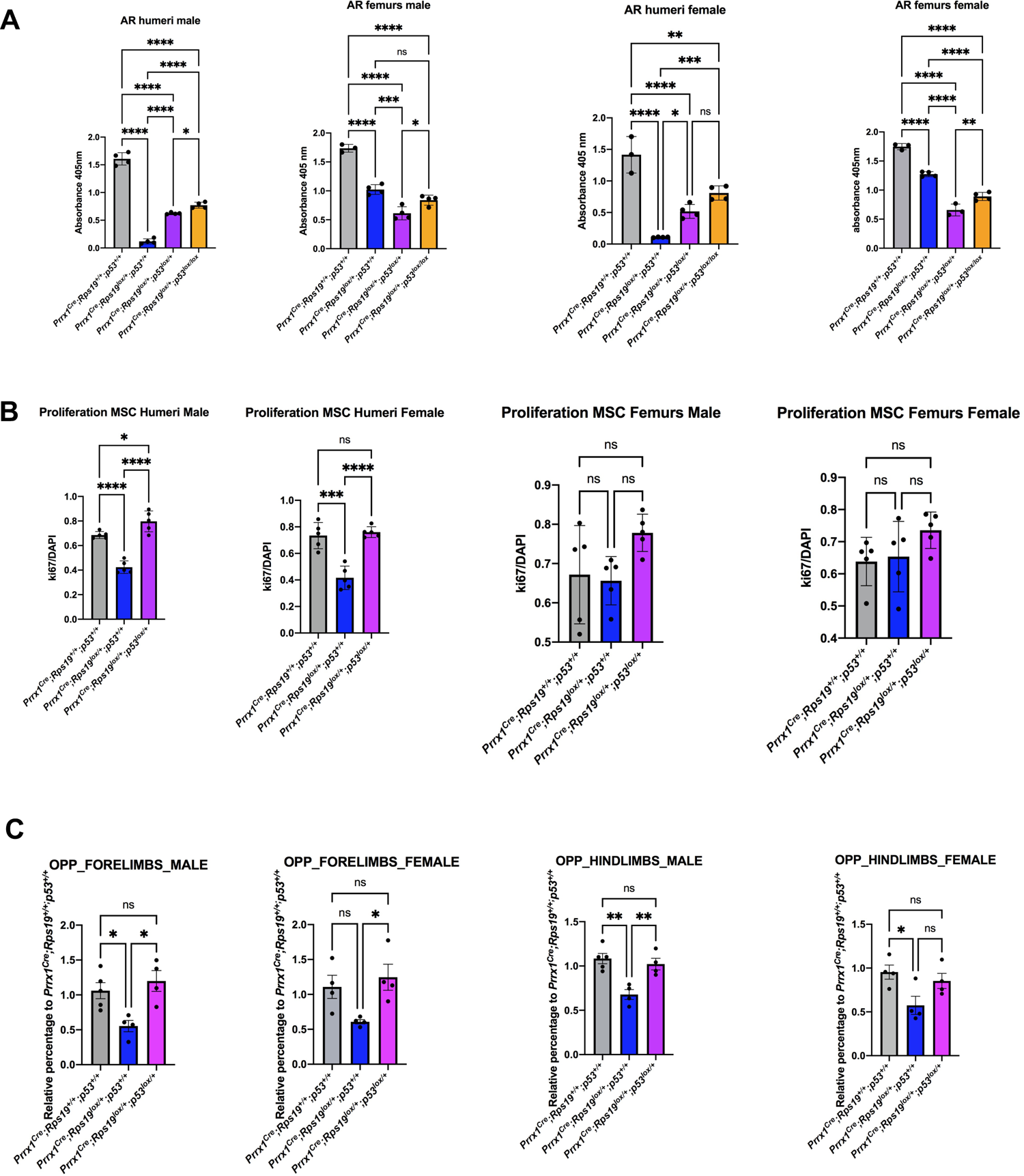
Sex specific differentiation, proliferation and protein synthesis phenotype in *Prrx1-Cre; Rps19^lox/+^* Mesenchymal Stem Cells. (A) Sex specific quantification of Alizarin Red S staining of whole 24-well plates wells of MSCs derived osteoblasts at day 14 of differentiation of *Prrx1-Cre; Rps19^+/+^, Prrx1-Cre; Rps19^lox/+^, Prrx1-Cre; Rps19^lox/+^p53^lox/+^*, *Prrx1-Cre; Rps19^lox/+^p53^lox/lox^* 3 - week - old mice. (n=4 for male *Prrx1-Cre; Rps19^+/+^*, n=4 for male *Prrx1-Cre; Rps19^lox/+^*, n=4 for male *Prrx1-Cre; Rps19^lox/+^p53^lox/+^*, n=4 for male *Prrx1-Cre; Rps19^lox/+^p53^lox/lox^* n=3 for female *Prrx1-Cre; Rps19^+/+^*, n=4 for female *Prrx1-Cre; Rps19^lox/+^*, n=4 for female *Prrx1-Cre; Rps19^lox/+^p53^lox/+^*, n=4 for female *Prrx1-Cre; Rps19^lox/+^p53^lox/lox^* (ANOVA, *p<0.05, **p<0.01, ***p<0.001). (B) Sex specific quantification of ki67 staining for MSCs of *Prrx1-Cre; Rps19^+/+^, Prrx1-Cre; Rps19^lox/+^, Prrx1-Cre; Rps19^lox/+^p53^lox/+^* 3 - week - old mice. (n=5 for male *Prrx1-Cre; Rps19^+/+^*, n=5 for male *Prrx1-Cre; Rps19^lox/+^*, n=5 for male *Prrx1-Cre; Rps19^lox/+^p53^lox/+^*, n=5 for female *Prrx1-Cre; Rps19^+/+^*, n=5 for female *Prrx1-Cre; Rps19^lox/+^*, n=5 for female *Prrx1-Cre; Rps19^lox/+^p53^lox/+^* (ANOVA, *p<0.05, **p<0.01, ***p<0.001). (C) Sex specific quantification for OPP incorporation in forelimb and hindlimb – derived MSCs of *Prrx1-Cre; Rps19^+/+^, Prrx1-Cre; Rps19^lox/+^, Prrx1-Cre; Rps19^lox/+^p53^lox/+^* 3 - week - old mice. (n=5 for male *Prrx1-Cre; Rps19^+/+^*, n=4 for male *Prrx1-Cre; Rps19^lox/+^*, n=4 for male *Prrx1-Cre; Rps19^lox/+^p53^lox/+^*, n=4 for female *Prrx1-Cre; Rps19^+/+^*, n=4 for female *Prrx1-Cre; Rps19^lox/+^*, n=4 for female *Prrx1-Cre; Rps19^lox/+^p53^lox/+^* ( (ANOVA, *p<0.05, **p<0.01, ***p<0.001).

**Supplemental Figure 6.**
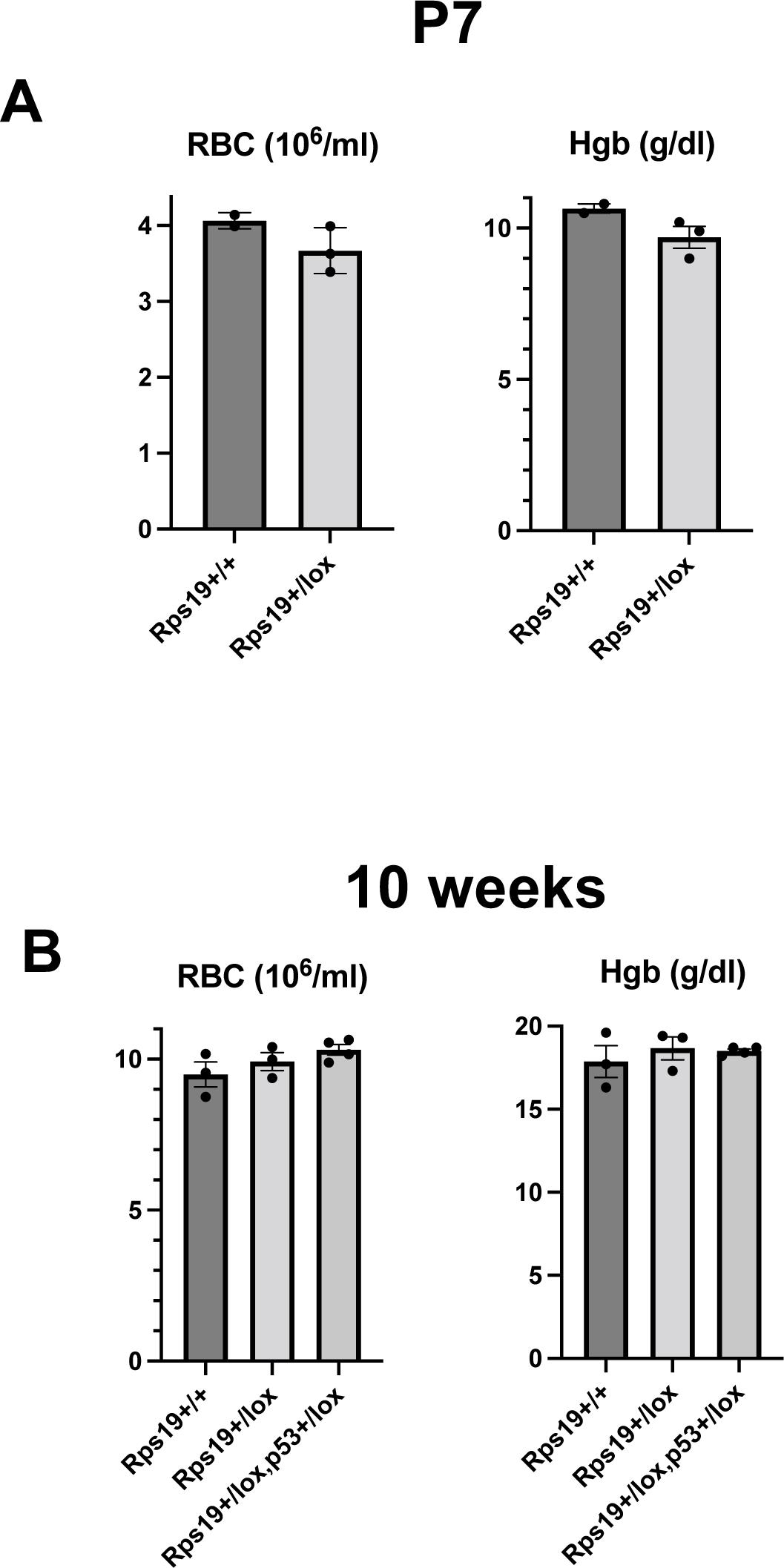
Red blood cell (RBC) counts and Hemoglobin (Hgb). (A) RBC and hemoglobin at day P7 in the *Prrx1-Cre; Rps19^+/+^* (black)*, Prrx1-Cre; Rps19^lox/+^* mice. (B) RBC and hemoglobin in 10 week old male *Prrx1-Cre; Rps19^+/+^, Prrx1-Cre; Rps19^lox/+^* and *Prrx1-Cre; Rps19^lox/+^p53^lox/+^* mice.

## Source Files

**Figure 2C-source data: uncropped gel of mESCs during osteoblast differentiation.** The arrow depicts Rps19 (page 1) and GAPDH (page 2). Lane number and correspondence presented on the side.

**Figure 3C-source data: uncropped gels of mESCs during osteoblast differentiation.** The arrow depicts Dvl3 (page 1), pLRP6 (page 2), LRP6 (page 3), β-catenin (page 4), β-catenin S552 (Page 5), β-catenin S675 (page 6), β-catenin S33/37/41 (page 7), GAPDH (page 8). Lane number and correspondence presented on the side.

**Figure 4C-source data: uncropped gels of RPS19 tissue expression.** The arrow depicts RPS19 and GAPDH in the various tissues and genotypes mentioned at the top.

**Figure 5B-source data: uncropped gel of mESCs during osteoblast differentiation**. The arrow depicts p53 (total) and GAPDH (bottom). Lane number and correspondence presented on the side.

**Figure 5C-source data: uncropped gel of mESCs during osteoblast differentiation**. The arrow depicts p53 (Ser15) and GAPDH (bottom). Lane number and correspondence presented on the side.

## References

1. Ambrosi, T. H., Marecic, O., McArdle, A., Sinha, R., Gulati, G. S., Tong, X., Wang, Y., Steininger, H. M., Hoover, M. Y., Koepke, L. S., Murphy, M. P., Sokol, J., Seo, E. Y., Tevlin, R., Lopez, M., Brewer, R. E., Mascharak, S., Lu, L., Ajanaku, O., … Chan, C. K. F. (2021). Aged skeletal stem cells generate an inflammatory degenerative niche. Nature, 597(7875), 256–262. https://doi.org/10.1038/s41586-021-03795-7

2. Artigas, N., Gamez, B., Cubillos-Rojas, M., Sanchez-de Diego, C., Valer, J. A., Pons, G., Rosa, J. L., & Ventura, F. (2017). p53 inhibits SP7/Osterix activity in the transcriptional program of osteoblast differentiation. Cell Death Differ, 24(12), 2022–2031. https://doi.org/10.1038/cdd.2017.113

3. Calvi, L. M., & Link, D. C. (2015). The hematopoietic stem cell niche in homeostasis and disease. Blood, 126(22), 2443–2451. https://doi.org/10.1182/blood-2015-07-533588

4. Da Costa, L., Chanoz-Poulard, G., Simansour, M., French, M., Bouvier, R., Prieur, F., Couque, N., Delezoide, A. L., Leblanc, T., Mohandas, N., & Touraine, R. (2013). First de novo mutation in RPS19 gene as the cause of hydrops fetalis in Diamond-Blackfan anemia. Am J Hematol, 88(2), 160. https://doi.org/10.1002/ajh.23366

5. Da Costa, L., Leblanc, T., & Mohandas, N. (2020). Diamond-Blackfan anemia. Blood, 136(11), 1262–1273. https://doi.org/10.1182/blood.2019000947

6. Danilova, N., & Gazda, H. T. (2015). Ribosomopathies: how a common root can cause a tree of pathologies. Dis Model Mech, 8(9), 1013–1026. https://doi.org/10.1242/dmm.020529

7. Doulatov, S., Vo, L. T., Macari, E. R., Wahlster, L., Kinney, M. A., Taylor, A. M., Barragan, J., Gupta, M., McGrath, K., Lee, H. Y., Humphries, J. M., DeVine, A., Narla, A., Alter, B. P., Beggs, A. H., Agarwal, S., Ebert, B. L., Gazda, H. T., Lodish, H. F., … Daley, G. Q. (2017). Drug discovery for Diamond-Blackfan anemia using reprogrammed hematopoietic progenitors. Sci Transl Med, 9(376). https://doi.org/10.1126/scitranslmed.aah5645

8. Duan, P., & Bonewald, L. F. (2016). The role of the wnt/beta-catenin signaling pathway in formation and maintenance of bone and teeth. Int J Biochem Cell Biol, 77(Pt A), 23–29. https://doi.org/10.1016/j.biocel.2016.05.015

9. Ducy, P., Zhang, R., Geoffroy, V., Ridall, A. L., & Karsenty, G. (1997). Osf2/Cbfa1: a transcriptional activator of osteoblast differentiation. Cell, 89(5), 747–754. https://doi.org/10.1016/s0092-8674(00)80257-3

10. Durland, J. L., Sferlazzo, M., Logan, M., & Burke, A. C. (2008). Visualizing the lateral somitic frontier in the Prx1Cre transgenic mouse. J Anat, 212(5), 590–602. https://doi.org/10.1111/j.1469-7580.2008.00879.x

11. Fumagalli, S., & Thomas, G. (2011). The role of p53 in ribosomopathies. Semin Hematol, 48(2), 97–105. https://doi.org/10.1053/j.seminhematol.2011.02.004

12. Gabut, M., Bourdelais, F., & Durand, S. (2020). Ribosome and Translational Control in Stem Cells. Cells, 9(2). https://doi.org/10.3390/cells9020497

13. Gay, D. M., Lund, A. H., & Jansson, M. D. (2021). Translational control through ribosome heterogeneity and functional specialization. Trends Biochem Sci. https://doi.org/10.1016/j.tibs.2021.07.001

14. Genuth, N. R., & Barna, M. (2018). The Discovery of Ribosome Heterogeneity and Its Implications for Gene Regulation and Organismal Life. Mol Cell, 71(3), 364–374. https://doi.org/10.1016/j.molcel.2018.07.018

15. Glatt, V., Canalis, E., Stadmeyer, L., & Bouxsein, M. L. (2007). Age-related changes in trabecular architecture differ in female and male C57BL/6J mice. J Bone Miner Res, 22(8), 1197–1207. https://doi.org/10.1359/jbmr.070507

16. Gray, J. D., Kholmanskikh, S., Castaldo, B. S., Hansler, A., Chung, H., Klotz, B., Singh, S., Brown, A. M., & Ross, M. E. (2013). LRP6 exerts non-canonical effects on Wnt signaling during neural tube closure. Hum Mol Genet, 22(21), 4267–4281. https://doi.org/10.1093/hmg/ddt277

17. Ishii, M., Egen, J. G., Klauschen, F., Meier-Schellersheim, M., Saeki, Y., Vacher, J., Proia, R. L., & Germain, R. N. (2009). Sphingosine-1-phosphate mobilizes osteoclast precursors and regulates bone homeostasis. Nature, 458(7237), 524–528. https://doi.org/10.1038/nature07713

18. Kanke, K., Masaki, H., Saito, T., Komiyama, Y., Hojo, H., Nakauchi, H., Lichtler, A. C., Takato, T., Chung, U. I., & Ohba, S. (2014). Stepwise differentiation of pluripotent stem cells into osteoblasts using four small molecules under serum-free and feeder-free conditions. Stem Cell Reports, 2(6), 751–760. https://doi.org/10.1016/j.stemcr.2014.04.016

19. Kim, N. H., Kim, H. S., Kim, N. G., Lee, I., Choi, H. S., Li, X. Y., Kang, S. E., Cha, S. Y., Ryu, J. K., Na, J. M., Park, C., Kim, K., Lee, S., Gumbiner, B. M., Yook, J. I., & Weiss, S. J. (2011). p53 and microRNA-34 are suppressors of canonical Wnt signaling. Sci Signal, 4(197), ra71. https://doi.org/10.1126/scisignal.2001744

20. Logan, M., Martin, J. F., Nagy, A., Lobe, C., Olson, E. N., & Tabin, C. J. (2002). Expression of Cre Recombinase in the developing mouse limb bud driven by a Prxl enhancer. Genesis, 33(2), 77–80. https://doi.org/10.1002/gene.10092

21. Lu, L., Harutyunyan, K., Jin, W., Wu, J., Yang, T., Chen, Y., Joeng, K. S., Bae, Y., Tao, J., Dawson, B. C., Jiang, M. M., Lee, B., & Wang, L. L. (2015). RECQL4 Regulates p53 Function In Vivo During Skeletogenesis. J Bone Miner Res, 30(6), 1077–1089. https://doi.org/10.1002/jbmr.2436

22. Ludwig, L. S., Gazda, H. T., Eng, J. C., Eichhorn, S. W., Thiru, P., Ghazvinian, R., George, T. I., Gotlib, J. R., Beggs, A. H., Sieff, C. A., Lodish, H. F., Lander, E. S., & Sankaran, V. G. (2014). Altered translation of GATA1 in Diamond-Blackfan anemia. Nat Med, 20(7), 748–753. https://doi.org/10.1038/nm.3557

23. Matsson, H., Davey, E. J., Draptchinskaia, N., Hamaguchi, I., Ooka, A., Leveen, P., Forsberg, E., Karlsson, S., & Dahl, N. (2004). Targeted disruption of the ribosomal protein S19 gene is lethal prior to implantation. Mol Cell Biol, 24(9), 4032–4037. https://doi.org/10.1128/MCB.24.9.4032-4037.2004

24. Mills, E. W., & Green, R. (2017). Ribosomopathies: There’s strength in numbers. Science, 358(6363). https://doi.org/10.1126/science.aan2755

25. Mortus, J. R., Zhang, Y., & Hughes, D. P. (2014). Developmental pathways hijacked by osteosarcoma. Adv Exp Med Biol, 804, 93–118. https://doi.org/10.1007/978-3-319-04843-7_5

26. Nakashima, K., Zhou, X., Kunkel, G., Zhang, Z., Deng, J. M., Behringer, R. R., & de Crombrugghe, B. (2002). The novel zinc finger-containing transcription factor osterix is required for osteoblast differentiation and bone formation. Cell, 108(1), 17–29. https://doi.org/10.1016/s0092-8674(01)00622-5

27. Narla, A., & Ebert, B. L. (2010). Ribosomopathies: human disorders of ribosome dysfunction. Blood, 115(16), 3196–3205. https://doi.org/10.1182/blood-2009-10-178129

28. Okamoto, M., Udagawa, N., Uehara, S., Maeda, K., Yamashita, T., Nakamichi, Y., Kato, H., Saito, N., Minami, Y., Takahashi, N., & Kobayashi, Y. (2014). Noncanonical Wnt5a enhances Wnt/beta-catenin signaling during osteoblastogenesis. Sci Rep, 4, 4493. https://doi.org/10.1038/srep04493

29. Olivares-Navarrete, R., Hyzy, S. L., Park, J. H., Dunn, G. R., Haithcock, D. A., Wasilewski, C. E., Boyan, B. D., & Schwartz, Z. (2011). Mediation of osteogenic differentiation of human mesenchymal stem cells on titanium surfaces by a Wnt-integrin feedback loop. Biomaterials, 32(27), 6399–6411. https://doi.org/10.1016/j.biomaterials.2011.05.036

30. Pederson, L., Ruan, M., Westendorf, J. J., Khosla, S., & Oursler, M. J. (2008). Regulation of bone formation by osteoclasts involves Wnt/BMP signaling and the chemokine sphingosine-1-phosphate. Proc Natl Acad Sci U S A, 105(52), 20764–20769. https://doi.org/10.1073/pnas.0805133106

31. Person, A. D., Beiraghi, S., Sieben, C. M., Hermanson, S., Neumann, A. N., Robu, M. E., Schleiffarth, J. R., Billington, C. J., Jr., van Bokhoven, H., Hoogeboom, J. M., Mazzeu, J. F., Petryk, A., Schimmenti, L. A., Brunner, H. G., Ekker, S. C., & Lohr, J. L. (2010). WNT5A mutations in patients with autosomal dominant Robinow syndrome. Dev Dyn, 239(1), 327–337. https://doi.org/10.1002/dvdy.22156

32. Phillips, M. D., Kuznetsov, S. A., Cherman, N., Park, K., Chen, K. G., McClendon, B. N., Hamilton, R. S., McKay, R. D., Chenoweth, J. G., Mallon, B. S., & Robey, P. G. (2014). Directed differentiation of human induced pluripotent stem cells toward bone and cartilage: in vitro versus in vivo assays. Stem Cells Transl Med, 3(7), 867–878. https://doi.org/10.5966/sctm.2013-0154

33. Rigueur, D., & Lyons, K. M. (2014). Whole-mount skeletal staining. Methods Mol Biol, 1130, 113–121. https://doi.org/10.1007/978-1-62703-989-5_9

34. Roifman, M., Marcelis, C. L., Paton, T., Marshall, C., Silver, R., Lohr, J. L., Yntema, H. G., Venselaar, H., Kayserili, H., van Bon, B., Seaward, G., Consortium, F. C., Brunner, H. G., & Chitayat, D. (2015). De novo WNT5A-associated autosomal dominant Robinow syndrome suggests specificity of genotype and phenotype. Clin Genet, 87(1), 34–41. https://doi.org/10.1111/cge.12401

35. Rowe, D. W., Adams, D. J., Hong, S. H., Zhang, C., Shin, D. G., Renata Rydzik, C., Chen, L., Wu, Z., Garland, G., Godfrey, D. A., Sundberg, J. P., & Ackert-Bicknell, C. (2018). Screening Gene Knockout Mice for Variation in Bone Mass: Analysis by muCT and Histomorphometry. Curr Osteoporos Rep, 16(2), 77–94. https://doi.org/10.1007/s11914-018-0421-4

36. Shi, Z., & Barna, M. (2015). Translating the genome in time and space: specialized ribosomes, RNA regulons, and RNA-binding proteins. Annu Rev Cell Dev Biol, 31, 31–54. https://doi.org/10.1146/annurev-cellbio-100814-125346

37. Sieff, C. (1993). Diamond-Blackfan Anemia. In M. P. Adam, H. H. Ardinger, R. A. Pagon, S. E. Wallace, L. J. H. Bean, G. Mirzaa, & A. Amemiya (Eds.), GeneReviews((R)). https://www.ncbi.nlm.nih.gov/pubmed/20301769

38. Sieff, C. A., Yang, J., Merida-Long, L. B., & Lodish, H. F. (2010). Pathogenesis of the erythroid failure in Diamond Blackfan anaemia. Br J Haematol, 148(4), 611–622. https://doi.org/10.1111/j.1365-2141.2009.07993.x

39. Singh, S. A., Goldberg, T. A., Henson, A. L., Husain-Krautter, S., Nihrane, A., Blanc, L., Ellis, S. R., Lipton, J. M., & Liu, J. M. (2014). p53-Independent cell cycle and erythroid differentiation defects in murine embryonic stem cells haploinsufficient for Diamond Blackfan anemia-proteins: RPS19 versus RPL5. PLoS One, 9(2), e89098. https://doi.org/10.1371/journal.pone.0089098

40. Teti, A., & Teitelbaum, S. L. (2019). Congenital disorders of bone and blood. Bone, 119, 71–81. https://doi.org/10.1016/j.bone.2018.03.002

41. Tiu, G. C., Kerr, C. H., Forester, C. M., Krishnarao, P. S., Rosenblatt, H. D., Raj, N., Lantz, T. C., Zhulyn, O., Bowen, M. E., Shokat, L., Attardi, L. D., Ruggero, D., & Barna, M. (2021). A p53-dependent translational program directs tissue-selective phenotypes in a model of ribosomopathies. Dev Cell, 56(14), 2089–2102 e2011. https://doi.org/10.1016/j.devcel.2021.06.013

42. Ulirsch, J. C., Verboon, J. M., Kazerounian, S., Guo, M. H., Yuan, D., Ludwig, L. S., Handsaker, R. E., Abdulhay, N. J., Fiorini, C., Genovese, G., Lim, E. T., Cheng, A., Cummings, B. B., Chao, K. R., Beggs, A. H., Genetti, C. A., Sieff, C. A., Newburger, P. E., Niewiadomska, E., … Gazda, H. T. (2018). The Genetic Landscape of Diamond-Blackfan Anemia. Am J Hum Genet, 103(6), 930–947. https://doi.org/10.1016/j.ajhg.2018.10.027

43. Vlachos, A., Rosenberg, P. S., Atsidaftos, E., Kang, J., Onel, K., Sharaf, R. N., Alter, B. P., & Lipton, J. M. (2018). Increased risk of colon cancer and osteogenic sarcoma in Diamond-Blackfan anemia. Blood, 132(20), 2205–2208. https://doi.org/10.1182/blood-2018-05-848937

44. Wang, Y., Li, Y. P., Paulson, C., Shao, J. Z., Zhang, X., Wu, M., & Chen, W. (2014). Wnt and the Wnt signaling pathway in bone development and disease. Front Biosci (Landmark Ed), 19, 379–407. https://doi.org/10.2741/4214

45. White, J. J., Mazzeu, J. F., Hoischen, A., Bayram, Y., Withers, M., Gezdirici, A., Kimonis, V., Steehouwer, M., Jhangiani, S. N., Muzny, D. M., Gibbs, R. A., Baylor-Hopkins Center for Mendelian, G., van Bon, B. W. M., Sutton, V. R., Lupski, J. R., Brunner, H. G., & Carvalho, C. M. B. (2016). DVL3 Alleles Resulting in a −1 Frameshift of the Last Exon Mediate Autosomal-Dominant Robinow Syndrome. Am J Hum Genet, 98(3), 553-561. https://doi.org/10.1016/j.ajhg.2016.01.005

46. Yu, V. W., & Scadden, D. T. (2016). Heterogeneity of the bone marrow niche. Curr Opin Hematol, 23(4), 331–338. https://doi.org/10.1097/MOH.0000000000000265

47. Zhou, X., Beilter, A., Xu, Z., Gao, R., Xiong, S., Paulucci-Holthauzen, A., Lozano, G., de Crombrugghe, B., & Gorlick, R. (2021). Wnt/ss-catenin-mediated p53 suppression is indispensable for osteogenesis of mesenchymal progenitor cells. Cell Death Dis, 12(6), 521. https://doi.org/10.1038/s41419-021-03758-w

48. Zhu, H., Guo, Z. K., Jiang, X. X., Li, H., Wang, X. Y., Yao, H. Y., Zhang, Y., & Mao, N. (2010). A protocol for isolation and culture of mesenchymal stem cells from mouse compact bone. Nat Protoc, 5(3), 550–560. https://doi.org/10.1038/nprot.2009.238

